# Prostaglandin D2 synthase controls Schwann cells metabolism

**DOI:** 10.1101/2024.02.29.582775

**Authors:** Amelia Trimarco, Matteo Audano, Rosa La Marca, Mariaconcetta Cariello, Marta Falco, Silvia Pedretti, Gabriele Imperato, Alessandro Cestaro, Paola Podini, Giorgia Dina, Angelo Quattrini, Luca Massimino, Donatella Caruso, Nico Mitro, Carla Taveggia

## Abstract

We previously reported that in the absence of Prostaglandin D2 synthase (L–PGDS) peripheral nerves are hypomyelinated in development and that with aging they present aberrant myelin sheaths. We now demonstrate that L–PGDS expressed in Schwann cells is part of a coordinated program aiming at preserving myelin integrity. *In vivo* and *in vitro* lipidomic, metabolomic and transcriptomic analyses confirmed that myelin lipids composition, Schwann cells energetic metabolism and key enzymes controlling these processes are altered in the absence of L–PGDS. Moreover, Schwann cells undergo a metabolic rewiring and turn to acetate as the main energetic source. Further, they produce ketone bodies to ensure glial cell and neuronal survival. Importantly, we demonstrate that all these changes correlate with morphological myelin alterations and describe the first physiological pathway implicated in preserving PNS myelin.

Collectively, we posit that myelin lipids serve as a reservoir to provide ketone bodies, which together with acetate represent the adaptive substrates Schwann cells can rely on to sustain the axo-glial unit and preserve the integrity of the PNS.

---

Maintenance of an intact communication between glial cells, Schwann cells in the peripheral nervous system (PNS) and oligodendrocytes in the central nervous system (CNS) is essential to ensure efficient and effective transmission of the nerve impulse. Myelin, the insulating plasma membrane made by myelinating Schwann cells in the PNS, is the main structure deputed to this function. In addition to facilitating the transmission of the electric impulses, myelin is also essential to preserve the communication between neurons and glial cells. Indeed, lack of myelin or alteration in its integrity results in permanent damage of the nervous system eventually leading to neuronal cell death ^1, 2^.

In the PNS, the interaction between glial cells and axons are composite and rely on an articulate set of molecules expressed both by Schwann cells and axons ^3^. Further, signaling coming from the extracellular matrix is also key to maintain the correct structure of the axo-glial unit ^4^. While the identity of these molecules and pathways have been defined in development and after nerve injury ^3, 5^, less is known on those implicated in the maintenance of this cross talk in adulthood to prevent myelin and/or axonal damage.

In addition, metabolism, and the transfer of key metabolites between glial cells and neurons are critical to maintain an intact nervous system. Indeed, either constitutive or cell specific ablation of enzymes regulating mitochondrial metabolism and energy, among which Cox10 ^6^, TFAM ^7, 8^ and LKB1^9^, confirmed that Schwann cells actively support axons. Further, it is becoming increasingly evident that not only glial cells’ metabolism uphold axons, but that both oligodendrocytes and Schwann cells transfer energy metabolites, essentially lactate, which is eventually used by neurons to sustain their energy demand ^6, 10^. In the PNS, it has been proposed that the excessive lactate production observed in LKB–1 deficient mutant Schwann cells could be transferred to neurons to limit axonal loss ^9^. However, specific ablation of the monocarboxylate transporter MCT1 in Schwann cells did not prevent myelination, though mutant mice have thinner myelin sheath in sensory nerves and impaired regeneration after injury ^11–14^. More recently it has been shown that immediately after injury, Schwann cells provide neurons with lactate to prevent accelerated axonal degeneration and promote the initiation of the nerve regeneration process ^15^.

We previously reported that prostaglandin D–2 synthase (L–PGDS) modulates PNS myelination and maintenance by acting downstream of Neuregulin 1 type III ^16^, the main instructive signal for PNS myelination^17^. In the absence of *l–pgds*, nerves are hypomyelinated during development, but in adulthood the structure of their myelin is aberrant. We now extend these studies by showing that these morphological alterations correlate with a switch specifically in Schwann cells’ metabolism. *In vivo* lipidomic analyses confirmed that the excessive myelin remodeling occurring in these mutants completely alters lipids’ profile. In particular levels of lysophosphatidylcholines containing the omega– 6 fatty acids, which are part of the Arachidonic acid metabolism, are significantly reduced. Of note, these modifications are accompanied by changes in Schwann cell energetic metabolism, which ultimately rely on acetate and ketone bodies to sustain glial cell and neuronal survival. Importantly, we observed a direct correlation between myelin instability and energy metabolism rewiring. Thus, our study indicates that similarly to the CNS ^18^ also in the PNS acetate metabolism could substitute for glycolytic activity.

To our knowledge this is the first study correlating prostaglandins to glial cell metabolism and myelin maintenance identifying acetate and ketone bodies utilization as key metabolic pathways in the PNS. Collectively, we posit that Schwann cells adapt their energetic demand to respond to excessive myelin remodeling by producing and resorting onto ketone bodies to sustain the axo-glial unit. In future studies, it could be of interest to investigate whether also in pathological conditions acetate is implemented to preserve nerve integrity and functionality.

## RESULTS

### Loss of l–pgds alters Schwann cell lipid homeostasis and energetic metabolism

We previously showed that in the absence of the PGD2 prostanoid, sciatic nerves are hypomyelinated and characterized by aberrant myelin in aged mutant mice lacking *h–* and *l–pgds* expression. We also reported that ablation of *l–pgds* alone is sufficient to cause hypomyelination ^16^. To define the molecular components at the basis of this altered myelin structure, we analyzed myelin protein expression as well as lipid and metabolism composition in sciatic nerves of null *l–pgds* and littermate controls at different ages. Quantitative RT–PCR (Fig. 1a–b, and 1d–e) and Western Blotting analyses (Fig. 1c and 1f) on sciatic nerves of 1– and 8– months old *l–pgds^-/-^* mice showed no differences in Myelin Basic Protein (MBP) and Protein Zero (P0) protein expression, indicating that myelin lipid profile rather than proteins might be affected in *l–pgds^-/-^* mice. Hence, we performed lipidomic and metabolomic analyses on sciatic nerves of 4–, 6– and 8–months old *l–pgds^-/-^* mutants. Principal component analyses (PCA) (Fig. 1g) and heat map analyses of the top 50 dysregulated metabolites, (Supplementary Fig. 1a– c) showed that both at 4 and at 6 months *l–pgds^-/-^* mice cluster separately from littermate controls, a difference less pronounced at 8 months (Fig. 1g), indicating that L–PGDS likely participates in lipid/metabolic homeostasis in sciatic nerves.

**Figure 1:**
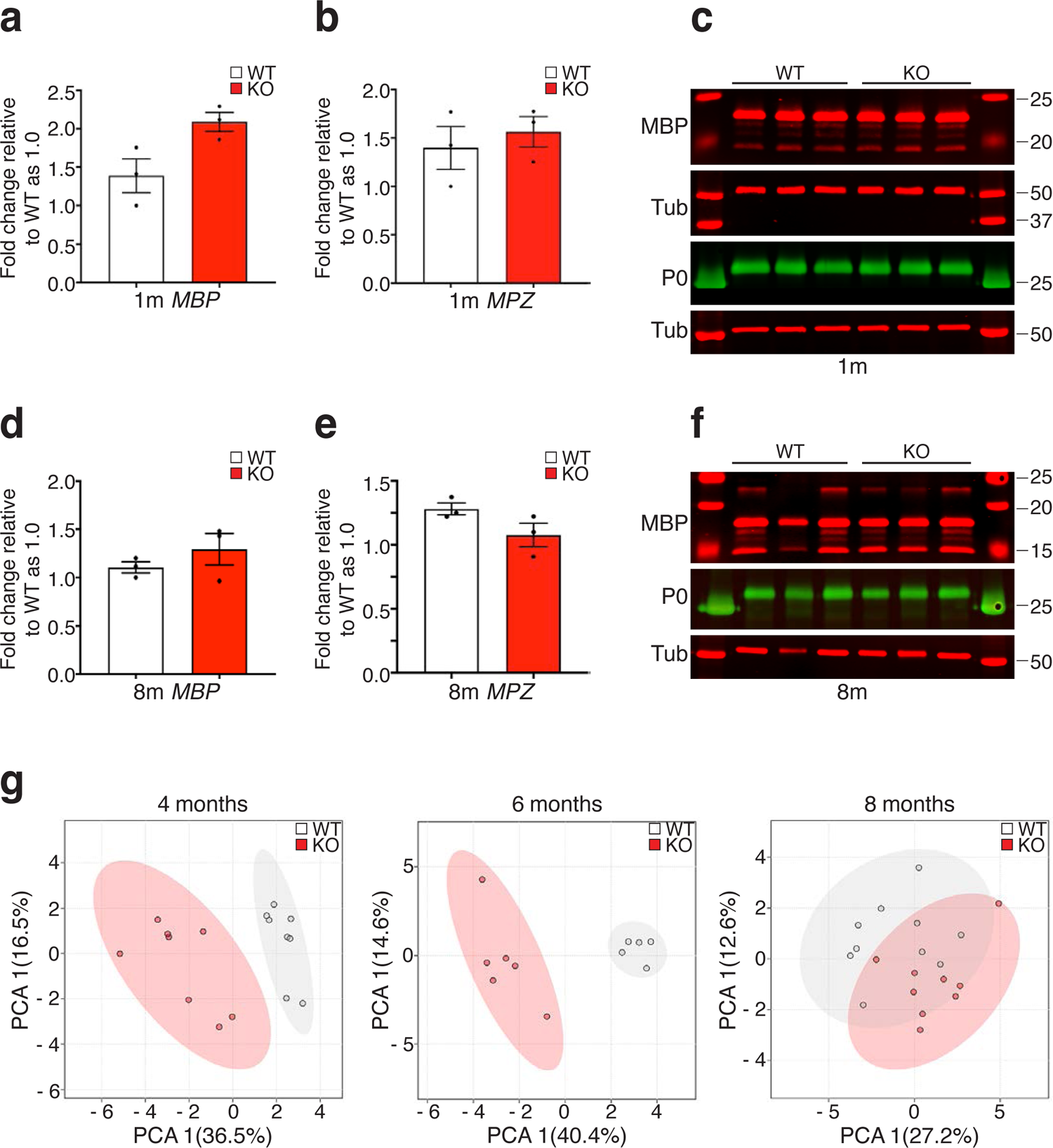
In the absence of L–PGDS Schwann cell lipid homeostasis and energetic metabolism are altered. **a**–**b**) qRT–PCR analyses on mRNA prepared from 1 month old wild type littermate controls (WT) and *l–pgds^-/-^* (KO) sciatic nerves and tested for *Mbp* (**a**) or *Mpz* (**b**) expression. While *Mbp* expression is slightly increased in null mutants, *Mpz* expression is comparable. Data have been normalized to *36B4* expression level and analyzed with the CFX Manager Software^TM^ from Biorad. N = 3 different mRNA preparations and analyses. Error bars represent mean ± s.e.m. (*Mbp* Unpaired t-test. P = 0.0502, t=2.773, df =4; *MPZ*: unpaired t test analysis, P = 0.5796; t=0.6020 df=4). **c**) Representative western blotting analyses on sciatic nerves prepared from 1 month old wild type littermate controls (WT) and *l–pgds^-/-^* (KO) sciatic nerves. Lysates were tested for MBP, P0 and Tubulin (Tub) as a loading control. **d**–**e**) qRT–PCR analyses on mRNA prepared from 8 months old wild type littermate controls (WT) and *l–pgds^-/-^*(KO) sciatic nerves and tested for *Mbp* (**a**) or *Mpz* (**b**) expression. No difference in the expression levels was observed between the genotypes. Data have been normalized to *36B4* expression level and analyzed with the CFX Manager Software^TM^ from Biorad. N = 3 different mRNA preparations and analyses. Error bars represent mean ± s.e.m. (*Mbp* Unpaired t-test. P = 0.346, t=1.067, df =4; *MPZ*: unpaired t test analysis, P = 0.1172; t=1.992 df=4) **f)** Representative western blotting analyses on sciatic nerves prepared from 8 months old wild type littermate controls (WT) and *l–pgds^-/-^* (KO) sciatic nerves. Lysates were tested for MBP, P0 and Tubulin (Tub) as a loading control. **g)** Graphs showing the Principal Component Analyses (PCA) of lipid and metabolites distribution in 4, 6 and 8 months old wild type littermate controls (WT) and *l–pgds^-/-^* (KO) sciatic nerves.

### Myelin alterations increase with age in sciatic nerves l–pgds^-/-^ mice

Next, we assessed whether these changes correlated with the previously reported altered morphology^16^. Thus, we quantified the number of myelin alterations in sciatic nerves in 4– and 8–months old *l–pgds^-/-^* mice and we observed that while at 8 months myelin aberrations are significantly increased in *l–pgds^-/-^* sciatic nerves, at 4 months, albeit augmented in mutant mice, they did not reach a significant difference (Fig. 2a–d). These *in vivo* results strongly suggest an important role for L–PGDS in myelin maintenance ^16^. Further, they indicate a trend in accumulating myelin alterations in adulthood.

**Figure 2.**
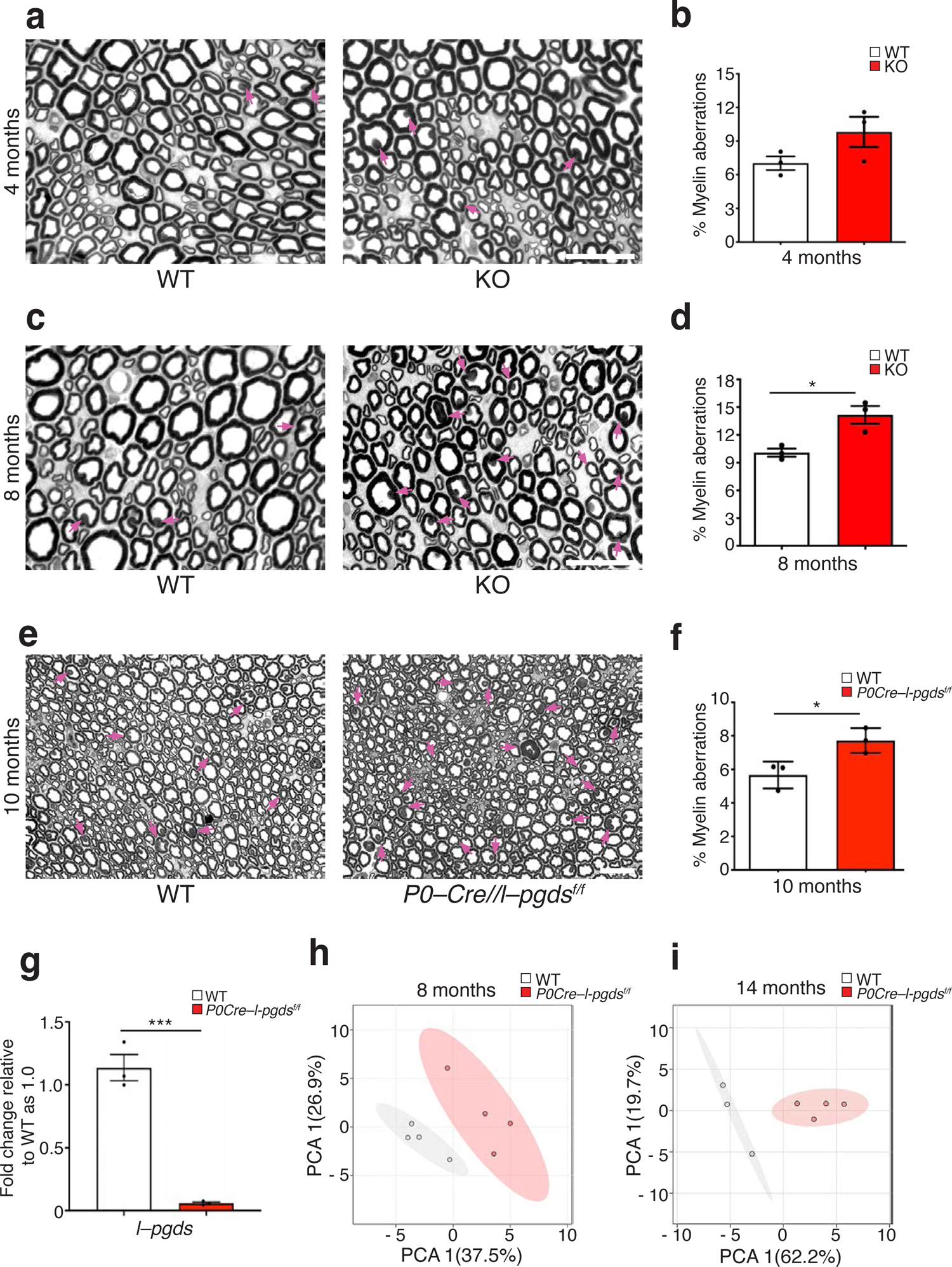
Glial L–PGDS controls myelin maintenance in aged mice. **a)** Semi–thin sections of 4 months wild–type littermate controls (WT) and *l–pgds^-/-^* sciatic nerves (KO). Bar: 50 μm. **b)** Graph, average of three different experiments, representing the percentage of myelin morphological aberrations in 4 months old wild type littermate controls (white bar) and *l–pgds^-/-^* sciatic nerves (red bar). Alterations were determined as the number of fibers presenting myelin structural alterations (pink arrows) over the total number of fibers in the entire reconstructed nerve cross section. Error bars represent mean ± s.e.m. N = 3 different mice/genotype. (Unpaired t test analysis, P = 0.1322; t=1.887 df=4). **c)** Semi–thin sections of 8 month wild–type littermate controls (WT) and *l–pgds^-/-^* sciatic nerves (KO). Bar: 50 μm. **d)** Graph, average of three different experiments, representing the percentage of myelin morphological aberrations in 8 months old wild type littermate controls (white bar) and *l–pgds^-/-^* sciatic nerves (red bar). Alterations were determined as the number of fibers presenting myelin structural alterations (pink arrows) over the total number of fibers in the entire reconstructed nerve cross section. Error bars represent mean ± s.e.m. N = 3 different mice/genotype. (Unpaired t test analysis, P = 0.0180; t=3.870 df=4). **e)** Semi–thin sections of 10 month wild–type littermate controls (WT) and *P0–Cre//l–pgds^flx/flx^*. Bar: 50 μm. **f)** Graph, average of three different experiments, representing the percentage of myelin morphological aberrations in 10 months old wild type littermate controls (white bar) and *P0–Cre//l–pgds^flx/flx^*sciatic nerves (red bar). Alterations were determined as the number of fibers presenting myelin structural alterations (pink arrows) over the total number of fibers in the entire reconstructed nerve cross section. Error bars represent mean ± s.e.m. N = 3 different mice/genotype. (Unpaired t test analysis, P = 0.0314; t=3.248 df=4). **g)** qRT–PCR analyses on mRNA prepared from P7 wild type littermate controls (WT) and *P0–Cre//l– pgds^flx/flx^* sciatic nerves tested for *l–pgds* expression. *l–pgds* is barely expressed in mutant nerves. Data have been normalized to *gapdh* expression level and analyzed with the CFX Manager Software^TM^ from Biorad. N = 3 different mRNA preparations and analyses. Error bars represent mean ± s.e.m. (Unpaired t-test. P = 0.0005, t=10.35, df =4). **h)** Graphs showing the Principal Component Analyses (PCA) of lipid and metabolites distribution in 8 months old wild type littermate controls (WT) and *P0–Cre//l–pgds^flx/flx^*sciatic nerves. **i)** Graphs showing the Principal Component Analyses (PCA) of lipid and metabolites distribution in 14 months old wild type littermate controls (WT) and *P0–Cre//l–pgds^flx/flx^* sciatic nerves.

### Myelin alterations accumulate in vivo in the absence of glial L–PGDS

Since L–PGDS is expressed both in neurons and in Schwann cells and neuronal L–PGDS contributes to myelin formation ^16^, we next sought to determine the cell specific effect of *l–pgds* in adulthood. Thus, we crossed *l–pgds^flx/flx^*mice ^19^ with mutant mice driving Cre recombinase expression either in motor neurons (*Chat–Cre*) ^20^ or in myelinating Schwann cells (*P0–Cre*) ^21^. We first confirmed specific recombination in ventral spinal cord (*Chat–Cre*) (Supplementary Fig. 2a) and in sciatic nerves (*P0–Cre*) (Fig. 2g). Ultrastructural analyses and *g* ratio measurements indicated that *Chat–Cre//l–pgds^flx/flx^* sciatic nerves were hypomyelinated at P7 (Supplementary Fig. 2b–e) but myelin was morphologically normal at 8 months (Supplementary Fig 2f–g), suggesting that neuronal L–PGDS might be implicated in regulating myelin formation, as formerly described ^16^. Next, we performed a detailed characterization of *P0–Cre//l–pgds^flx/flx^* sciatic nerves at different time points, from P7 to adulthood (Fig. 2e-f and Supplementary Fig. 3). Morphological analyses and *g* ratio measurements indicated that, unlike *Chat– Cre//l–pgds^flx/flx^* mutants, myelin formation was normal in the absence of glial *l-pgds*, as confirmed by ultrastructural analyses at P7 and 1 month (Supplementary Fig. 3). However, we observed an increase in the number of myelin alterations, primarily evident from 10 months old *P0–Cre//l–pgds^flx/flx^* mice (Fig. 2e–f), as in complete null *l–pgds* mice. Based on these observations, we performed metabolomic/lipidomic profile in *P0–Cre//l–pgds^flx/flx^*mice before (8 months) and after the appearance of these alterations (14 months) to determine whether, as suggested by morphological analyses, they clustered differently from controls. Indeed, PCA (Fig. 2h–i) and heat map analyses of the top 50 dysregulated metabolites, (Supplementary Fig. 1d–e) showed that, similarly to *l–pgds^-/-^* mutants, *P0– Cre//l–pgds^flx/flx^* mice clustered separately from littermate controls already at 8 months, before the accumulation of aberrant myelin profiles and, even more strikingly at 14 months, suggesting that glial L–PGDS could regulate lipid/metabolic homeostasis in sciatic nerves.

Collectively, these *in vivo* analyses strongly implicate glial L–PGDS in controlling myelin homeostasis ^16^. Further, they also indicate that in the PNS glial and neuronal L–PGDS exerts different roles, with the latter being more important for myelin formation.

### Peripheral nerve lipid metabolism is impaired in the absence of glial l–pgds^-/-^

Since L–PGDS is an enzyme of Arachidonic acid metabolism, which, together with omega–6 fatty acids, is naturally present in myelin phospholipids, we retrieved phospholipid analyses on total extracts of 4–, 6– and 8–months old *l–pgds^-/-^* sciatic nerves as well as in 8–, 10– and 14–months *P0–Cre//l– pgds^flx/flx^*old mice, to better characterize the differences observed in PCA analyses.

Interestingly, despite the PCA analyses, phospholipids’ profile was similar in 4–months old *l–pgds^-/-^* and control mice (Fig. 3a). However, at 6 (Fig. 3b) and at 8–months of age (Fig. 3c), *l–pgds^-/-^*sciatic nerves were mostly altered as compared to littermate wild type controls. We observed a strong significant decrease (p<0.001) in lysophosphatidylcholines containing Arachidonic acid (LPC 20:4) and in omega– 6 fatty acids precursors of Arachidonic acid, such as Linoleic acid (LPC 18:2) and di–homo–γ linoleic acid (LPC 20:3) (Supplementary Fig. 4a–d). Notably, in 8 months old *l–pgds^-/-^* sciatic nerves we also observed a decrease in both total and free cholesterol, a critical component of myelin, whose amounts were not affected at 4 months (Fig. 3d–e).

**Figure 3:**
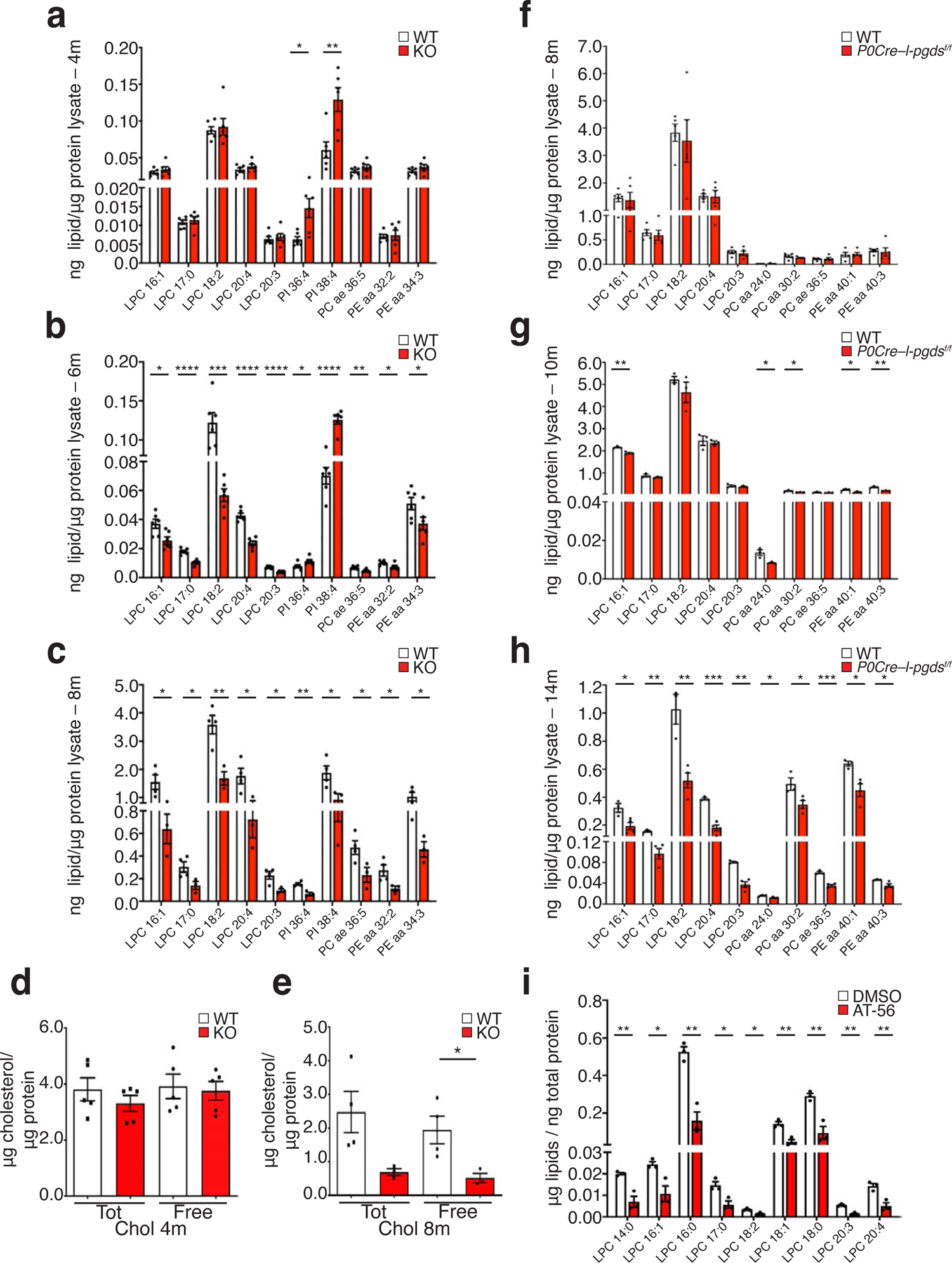
Targeted lipidomic analyses show alteration in phospholipids’ profile in the absence of l–pgds. **a)** Graph showing phospholipids profile in sciatic nerves of 4 months old wild type littermate controls (WT) and *l–pgds^-/-^* (KO) mice. Lipids were normalized over the total protein content. Error bars represent mean ± s.e.m. N = 6 different mice/genotype. (Unpaired t-test * P = 0.0105; t=3.141, df =10, **P = 0.0057; t=3.505, df=10). **b)** Graph showing phospholipids profile in sciatic nerves of 6 months old wild type littermate controls (WT) and *l–pgds^-/-^* (KO) mice. Lipids were normalized over the total protein content. Error bars represent mean ± s.e.m. N = 5 different mice/genotype. (Unpaired t-test. LPC 16:1 *P = 0.0157, t=2.906, df =10; LPC 17:0 ****P <0.0001, t=7.231, df =10; LPC 18:2 ***P =0.0007 t=4.867, df =10; LPC 20:4 ****P <0.0001, t=8.631, df =10; LPC 20:3 ****P <0.0001, t=8.214, df =10; PI 36:4 *P = 0.0294, t=2.54, df =10; PI 38:4 ****P <0.0001, t=7.169, df =10; PC ae 36:5 **P =0.0039 t=3.739, df =10; PE aa 32:2 *P = 0.029, t=2.547, df =10; PE aa *P = 0.0484, t=2.247, df =10). **c)** Graph showing phospholipids profile in sciatic nerves of 8 months old wild type littermate controls (WT) and *l–pgds^-/-^* (KO) mice. Lipids were normalized over the total protein content. N = 4 wild type and N = 3 null mice. Error bars represent mean ± s.e.m. (Unpaired t-test. LPC 16:1 *P = 0.0373, t=2.815, df =5; LPC 17:0 *P = 0.0438, t=2.68, df =5; LPC 18:2 **P = 0.0074 t=4.346, df =5; LPC 20:4 *P = 0.0314, t=2.962, df =5; LPC 20:3 *P = 0.0197, t=3.38, df =5; PI 36:4 **P = 0.0036, t=5.159, df =5; PI 38:4 *P = 0.0396, t=2.764, df =5; PC ae 36:5 *P =0.0434 t=2.688, df =5; PE aa 32:2 *P = 0.0452, t=2.653, df =5; PE aa *P = 0.0314, t=2.963, df =5). **d)** Graph showing total and free cholesterol levels in sciatic nerves of 4 months old wild type littermate controls (WT) and *l–pgds^-/-^* (KO) mice. Cholesterol levels were normalized over the total protein content. Variation in cholesterol amount is expressed as fold change to WT arbitrarily set as 1.0. N = 5 different mice/genotype. Error bars represent mean ± s.e.m. (Unpaired t-test. Tot: P = not significant, t=1.003, df =8; Free: P = not significant, t=0.292, df =8). **e)** Graph showing total and free cholesterol levels in sciatic nerves of 8 months old wild type littermate controls (WT) and *l–pgds^-/-^* (KO) mice. Cholesterol levels were normalized over the total protein content. Variation in cholesterol amount is expressed as fold change to WT arbitrarily set as 1.0. N = 4 wild type and N= 3 null mice. Error bars represent mean ± s.e.m. (Unpaired t-test. Tot: P = 0.0588, t=2.438, df =5; Free: P = 0.0346, t=2.88, df =5). **f)** Graph showing phospholipids profile in sciatic nerves of 8 months old wild type littermate controls (WT) and *P0–Cre//l–pgds^flx/flx^*mice. Lipids were normalized over the total protein content. Error bars represent mean ± s.e.m. N = 3 different mice/genotype. (Unpaired t test. No significant difference in any of the analyzed metabolites). **g)** Graph showing phospholipids profile sciatic nerves of 10 months old wild type littermate controls (WT) and *P0–Cre//l–pgds^flx/flx^* mice. Lipids were normalized over the total protein content. Error bars represent mean ± s.e.m. N = 5 different mice/genotype. (Unpaired t-test. LPC 16:1 **P = 0.0072, t=5.066, df =4; PC 24:0 *P = 0.0232, t=3.578, df =4; PC 30:2 *P = 0.0162, t=3.998, df =4; PE 40:1 *P = 0.028, t=3.372, df =4; PE 40:3 **P = 0.0037, t=6.092, df =4) **h)** Graph showing phospholipids profile in sciatic nerves of 14 months old wild type littermate controls (WT) and *P0–Cre//l–pgds^flx/flx^*mice. Lipids were normalized over the total protein content. Error bars represent mean ± s.e.m. N = 4 *P0–Cre//l–pgds^flx/flx^* mice; / N = 3 littermate controls (WT) mice. (Unpaired t-test. LPC 16:1 *P = 0.0248, t=3.171, df =5; LPC 17:0 **P = 0.0055, t=4.665, df =5; LPC 18:2 **P = 0.0053, t=4.697, df =5; LPC 20:4 ***P = 0.0005, t=8.018, df =5; LPC 20:3 **P = 0.0012, t=6.628, df =5; PC 24:0 *P = 0.0214, t=3.305, df =5; PC 30:2 *P = 0.0341, t=2.891, df =5; PC 36:5 ***P = 0.0004, t=8.208, df =5; PE 40:1 *P = 0.0173, t=3.501, df =5; PE 40:3 *P = 0.0351, t=2.868, df =5). **i)** Graph showing the phospholipids profile in Schwann cell – neuronal myelinated cocultures. After 14 days in myelinating conditions, cultures were treated for additional 7 days with 25 μM AT–56 or with DMSO as control vehicle. Levels of lysophosphatydilcholines containing omega–6 fatty acids are reduced in myelinated cocultures treated with AT–56. Lipids were normalized over the total protein content. N = 3 different independent coculture experiments. Error bars represent mean ± s.e.m. (Unpaired t-test. LPC 14:0 **P = 0.0073, t=5.035, df =4; LPC 16:1 *P = 0.0248, t=3.503, df =4; LPC 16:0 **P = 0.0026, t=6.716, df =4; LPC 17:0 *P = 0.0188, t=3.818, df =54; LPC 18:2 *P = 0.0132 t=4.243, df =4; LPC 18:1 **P = 0.0055 t=5.447, df =4; LPC 18:0 **P = 0.006 t=5.316, df =4; LPC 20:3 **P = 0.0011, t=8.485, df =4; LPC 20:4 **P = 0.0086, t=4.802, df =4).

Targeted lipidomic analyses in *P0–Cre//l–pgds^flx/flx^*sciatic nerves revealed similar alterations in phospholipid composition (Fig. 3f–h), which were mostly significant in older mice. Thus, although we cannot exclude that L–PGDS regulates plasma membrane lipid composition in several cell types, the striking reduction observed in at 14 months *P0–Cre//l–pgds^flx/flx^* sciatic nerves coupled to the myelin morphological alterations strongly impinge on a central role of glial L–PGDS in controlling lipid metabolism in peripheral nerves.

To further corroborate these results, we also performed targeted phospholipid analyses on organotypic wild–type myelinated DRG – Schwann cells cocultures treated with 25 μM AT–56, a specific irreversible L–PGDS inhibitor ^22^, for 7 days. Similarly to our *in vivo* results, we observed a significant decrease in lysophosphatidylcholines containing Arachidonic acid and in omega–6 fatty acids specifically in cocultures treated with AT–56, namely LPC 20:4, LPC 18:2 and LPC 20:3 (Fig. 3i). Globally, these data indicate that in the absence of L–PGDS lipid metabolism is remarkably altered in Schwann cells.

### Loss of l–pgds in sciatic nerves from aged mice decreases glycolysis and Krebs cycle activity

The PCA analyses reported in Fig. 1 and in Fig. 2, suggest that not only lipid metabolism is altered in the absence of *l–pgds*, but that also the energetic metabolism is affected in sciatic nerves of *l–pgds^-/-^* and of *P0–Cre//l–pgds^flx/flx^*mice. Thus, we examined the data relative to the energetic metabolites from sciatic nerves of both mutants collected at 4–, 6– and 8– months (*l–pgds^-/-^)* and at 8–, 10– and 14– months (*P0–Cre//l–pgds^flx/flx^*).

While at 4–months, in *l–pgds^-/-^* sciatic nerves we did not observe any significant alteration (Fig. 4a), both at 6– (Fig. 4b) and at 8–months (Fig. 4c), metabolites involved in glycolysis and Krebs cycle were overall reduced, and, as in the case of phospholipids, this effect was more significant at 6 months. Indeed, at this age we detected an increase in glucose–6P/fructose–6P accompanied by a decrease in glyceraldehyde–3–phosphate/dihydroxyacetone–phosphate (DHAP/GAP) and pyruvate that are indicative of an overall slowdown in glycolysis (Fig. 4b). Further, the analyses of the Krebs cycle revealed a reduction in citrate, succinyl–CoA, fumarate, malate and oxaloacetate in *l–pgds^-/-^* nerves. Notably, we observed a decrease also in acetyl–CoA in 8 months old mice (Fig. 4c and Supplementary Fig. 4e). Since citrate and α–ketoglutarate (Fig. 4c and Supplementary Fig. 4f–g) were also reduced in 8 months old sciatic nerves, these data suggest that upon loss of L–PGDS, there is a change in acetyl–CoA metabolic production/utilization in the Krebs cycle featuring a metabolic rewiring occurring in Schwann cells.

**Figure 4:**
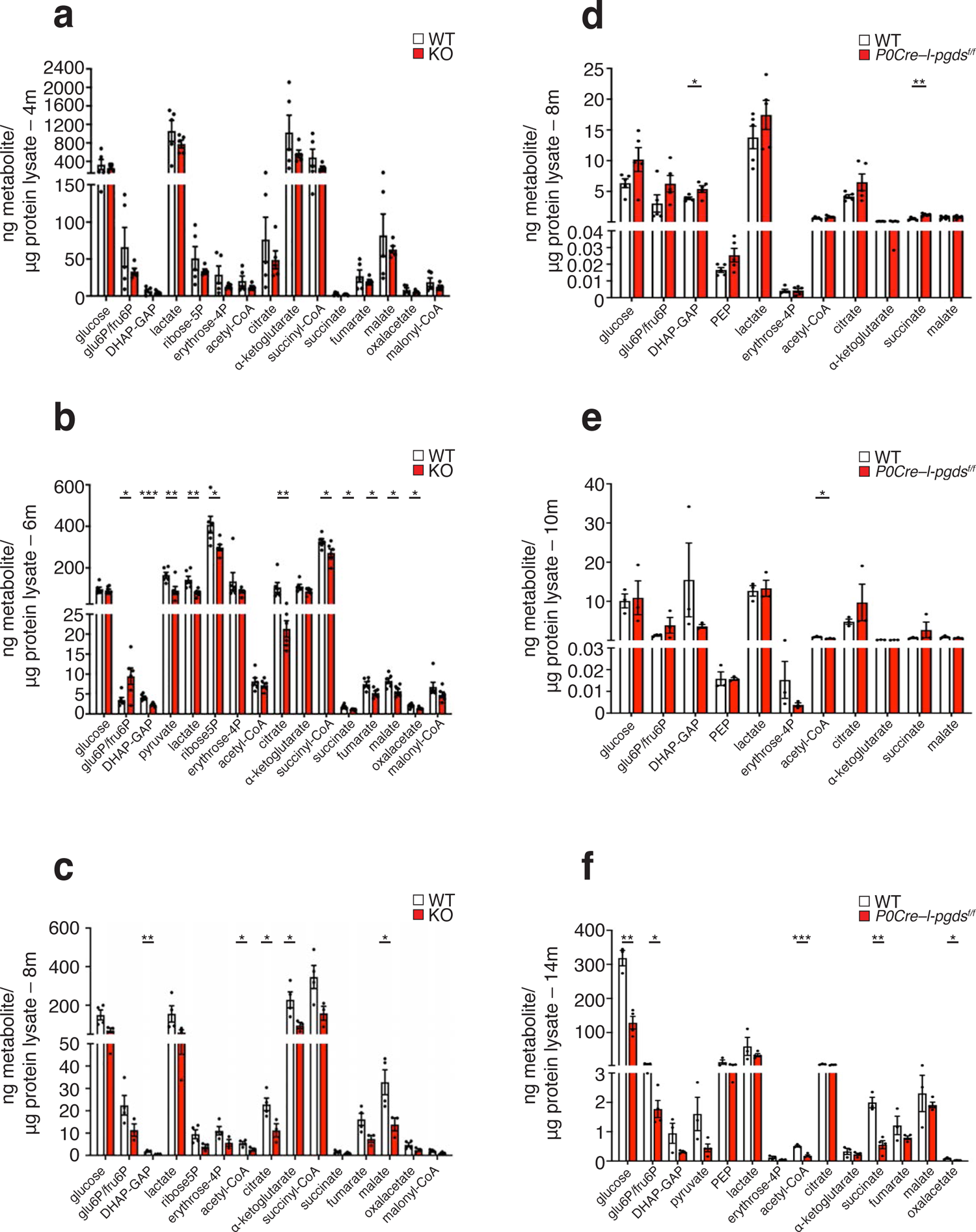
L–PGDS controls glycolysis and Krebs cycle metabolites in sciatic nerves. **a)** Graph showing the metabolic profile in 4 months old wild type littermate controls (WT) and *l–pgds^-/-^* (KO) sciatic nerves. Metabolites were normalized over the total protein content. N = 5 different mice/genotype. Error bars represent mean ± s.e.m. (Unpaired t-test. No significant difference in any of the analyzed metabolites). **b)** Graph showing the metabolic profile in 6 months old wild type littermate controls (WT) and *l–pgds^-/-^* (KO) sciatic nerves. Metabolites were normalized over the total protein content. N = 6 different mice/genotype. Error bars represent mean ± s.e.m. (Unpaired t-test. Glu6P/fru6P *P = 0.0196, t=2.774, df =10; DHAP–GAP ***P = 0.0004, t=5.264, df =10; pyruvate **P = 0.0098, t=3.178, df =10; lactate **P = 0.0066, t=3.418, df =10; ribose5P *P = 0.0272, t=2.585, df =10; citrate **P = 0.0047, t=3.614, df =10; succinyl–CoA *P = 0.0348, t=2.441, df =10; succinate *P = 0.0121, t=3.057, df =10; fumarate *P = 0.0136, t=2.988, df =10; malate *P = 0.0133, t=3.001, df =10; oxalacetate *P = 0.0194, t=2.78, df =10). **c)** Graph showing the metabolic profile in 8 months old wild type littermate controls (WT) and *l–pgds^-/-^* (KO) sciatic nerves. Metabolites were normalized over the total protein content. N = 4 wild type and N = 3 null mice. Error bars represent mean ± s.e.m. (Unpaired t-test. DHAP–GAP **P = 0.0071, t=4.392, df =5; acetyl–CoA *P = 0.0367, t=2.828, df =5; citrate *P = 0.038, t=2.799, df =5; α–ketoglutarate *P = 0.0473, t=2.617, df =5; malate *P = 0.0433, t=2.689, df =5). **d)** Graph showing the metabolic profile in 8 months old wild type littermate controls (WT) and *P0– Cre//l–pgds^flx/flx^* sciatic nerves. Metabolites were normalized over the total protein content. N = 3 different mice/genotype. Error bars represent mean ± s.e.m. (Unpaired t-test. DHAP–GAP *P = 0.0301, t=2.631, df =8; succinate **P = 0.0018, t=4.58, df =8). **e)** Graph showing the metabolic profile in 10 months old wild type littermate controls (WT) and *P0– Cre//l–pgds^flx/flx^* sciatic nerves. Metabolites were normalized over the total protein content. N = 5 different mice/genotype. Error bars represent mean ± s.e.m. (Unpaired t-test. Acetyl–CoA *P = 0.0221, t=3.632, df =4). **e**) Graph showing the metabolic profile in 14 months old wild type littermate controls (WT) and *P0– Cre//l–pgds^flx/flx^*sciatic nerves. Metabolites were normalized over the total protein content. N = 4 *P0– Cre//l–pgds^flx/flx^* mice; / N = 3 littermate controls (WT) mice. Error bars represent mean ± s.e.m. (Unpaired t-test. Glucose **P = 0.0013, t=6.532, df =5; Glu6P/fru6P *P = 0.028, t=3.063, df =5; Acetyl– CoA ***P = 0.0009, t=7.035, df =5; succinate **P = 0.0013, t=6.475, df =5; oxalacetate *P = 0.014, t=3.702, df =5).

Previous studies have shown that both oligodendrocytes and Schwann cells, can convert pyruvate in lactate, which is then transferred to neurons via monocarboxylate transporters ^2, 10, 15, 23, 24^. To determine whether in the absence of L–PGDS, glucose is converted to lactate, we measured this metabolite in 4–, 6– and 8–months old sciatic nerves of *l–pgds^-/-^* mutants and littermate controls. To our surprise, both at 6– and 8–months of age, lactate levels were reduced in mutant mice relative to controls (Fig. 4b–c and Supplementary Fig. 4h).

Surprisingly, metabolites’ changes in *P0–Cre//l–pgds^flx/flx^* mice at all analyzed time points (Fig. 3d– f), were less pronounced than in complete null mice. Nevertheless, we observed a significant reduction in glucose and DHAP/GAP, suggestive also in this case of a slowdown in glycolysis. Strikingly, levels of acetyl–CoA were also strongly decreased at 14 months of age, along with those of succinate and oxaloacetate, indicating a relevant impairment in Schwann cells Krebs cycle. Nevertheless, these cells did not experience energy depletion (Fig. 3f).

Collectively, these results indicate that in the absence of *l–pgds* both lipidomic and energetic profiles are altered. Though these changes are more evident in the complete *l–pgds* null sciatic nerves, aged mice lacking *l–pgds* in Schwann cells have an important reduction in glycolysis and Krebs cycle, thus suggesting that these glial cells may rely on alternative metabolic pathways.

### In the absence of l–pgds genes involved in lipid homeostasis and cell metabolism are upregulated

To characterize the molecular mechanism at the basis of these morphological alterations, we performed RNAseq analyses on wild–type myelinated mouse dorsal root ganglia (DRG) neurons – rat Schwann cells cocultures treated with 25 μM AT–56, a specific irreversible inhibitor of L–PGDS ^22^. As previously reported ^16^, we observed myelin degeneration reproducing the *in vivo* phenotype. To exclude that the myelin phenotype observed in AT–56 treated cocultures, might be due to Schwann cells and/or neuronal cell death we performed TUNEL assays analyses on myelinated cocultures treated with AT– 56, which, in agreement with our former studies ^16^, did not reveal any cell death in either cell types, or axonal swallowing (Supplementary Fig. 5a).

Next, we prepared and sequenced RNAs from cocultures treated with AT–56 versus vehicle only treated cocultures, as control. Overall, the number of upregulated genes significantly outnumbered downregulated ones. To determine whether identified genes belonged to neurons (mouse) or Schwann cells (rat), we aligned identified sequences to both mouse and rat deposited exome sequences (derived from NCBI assembly databases Build 37/mm9 and RGSC_v3.4 respectively) and selected genes with a P value < 0.001. From the alignment on the mouse exome sequence, we detected a total of 30 differentially expressed genes, 18 of which were upregulated (FC > 1.7) and 12 downregulated genes (FC < 0.5). Similarly, from the alignment on rat exome sequence we identified a total of 281 genes, 24 of which upregulated genes (FC > 1.7) and 2 downregulated (FC < 0.5). Of note, only rat genes, whose identity differs from those aligning to the mouse sequence, showed functional relationships, indicating that L–PGDS inhibition alters gene expression mainly in Schwann cells (Fig. 5a).

**Figure 5:**
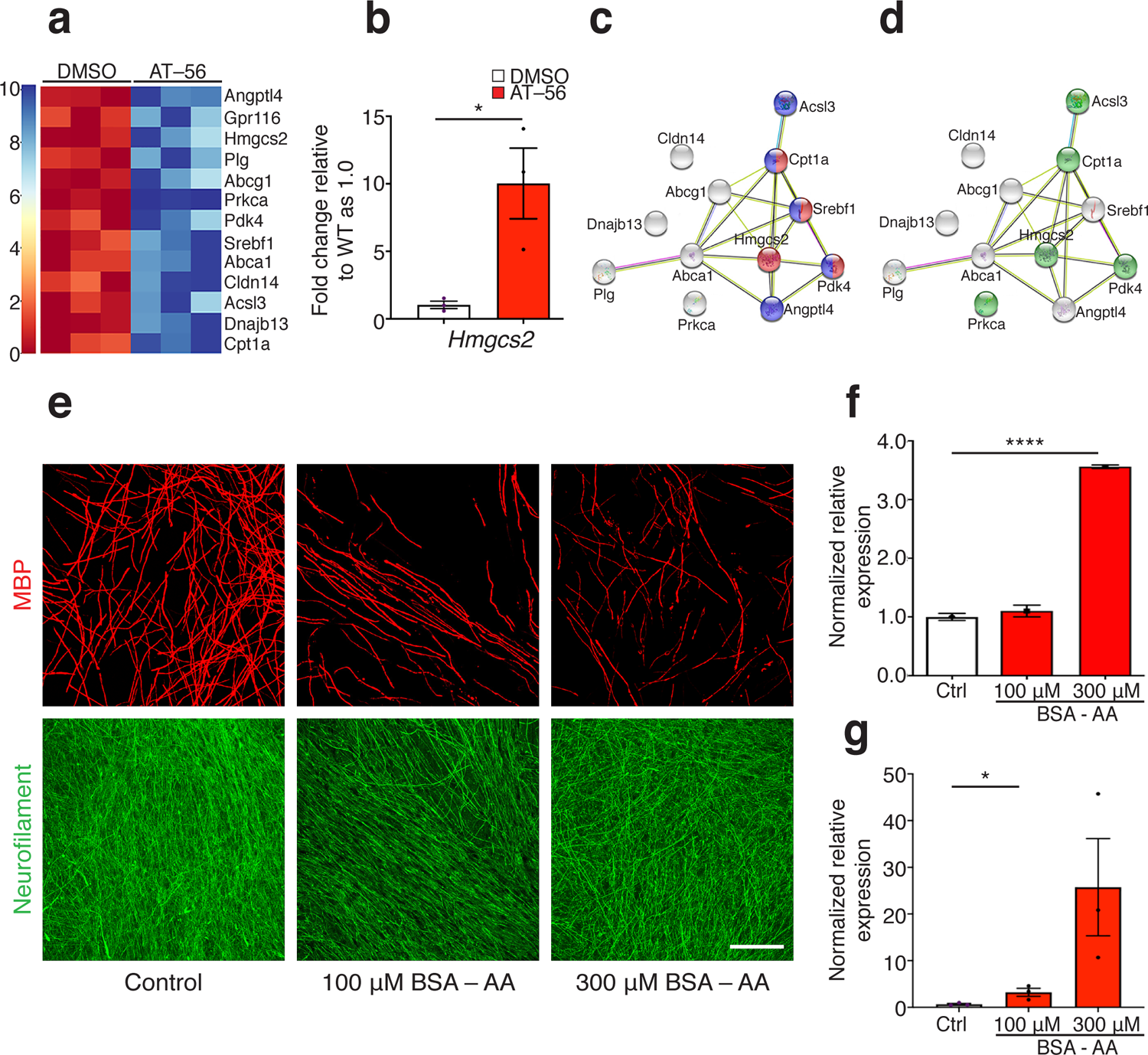
L–PGDS regulates genes involved in lipid homeostasis and Schwann cell energetic metabolism. **a)** Heat–map showing the main upregulated genes obtained with RNAseq analysis in rat primary Schwann cells – mouse neuronal myelinating cocultures. After 14 days in myelinating conditions, cultures were treated for additional 7 days with 25 μM AT–56 or with DMSO as control vehicle before processing. (Alignments on rat genome with P < 0.001). **b)** qRT–PCR analyses on mRNA prepared from rat Schwann cell – mouse neuronal cocultures myelinated for 14 days and then treated for additional 7 days with 25 μM AT–56 or with DMSO as vehicle control. The expression of *Pdk4*, *Hmgcs2*, *Acsl3* and *Angptl4* is increased in AT–56 treated cocultures. Data have been normalized to *gapdh* expression level and analyzed with the StepOne Software v2.3 for *Pdk4*, *Acsl3* and *Angptl4* (Applied Biosystems). The expression of *Hmgcs2* has been normalized to *TBP* expression level and analyzed with the CFX Manager Software^TM^ from Biorad. N = 3 different mRNA preparations and analyses. Error bars represent mean ± s.e.m. Error bars represent mean ± s.e.m. (Two–way ANOVA; *pdk4* ****P < 0.0001 F (2, 12) = 256.5; *Hmgcs2*: unpaired t-test. *P = 0.0268, t=3.418, df =4; *Acsl3* ***P = 0.0004 F F (2, 12) =15.92; *Angptl4* ****P < 0.0001 F (2, 12) = 315.1). **c)** qRT–PCR analyses on mRNA prepared from 10 months old wild type littermate controls (WT) and *P0–Cre//l–pgds^flx/flx^* sciatic nerves tested for P*dk4* and *Hmgcs2* expression. Data have been normalized to *36B4* expression level and analyzed with the CFX Manager Software^TM^ from Biorad. The expression of both genes is increased in *P0–Cre//l–pgds^flx/flx^* sciatic nerves. N = 4 (*Pdk4*) N = 6 (*Hmgcs2*) different mRNA preparations and analyses. Error bars represent mean ± s.e.m. (Unpaired t-test. *Pdk4* *P = 0.0308, t=2.808, df =6; *Hmgcs2*: *P = 0.0213, t=2.854, df =8). **d)** STRING pathway analyses and functional enrichments identified genes involved in “cellular response to fatty acid” (GO:0071398) and in “regulation of lipid metabolic process” (GO:0019216) with a false discovery rate of 2.13e–05. **e)** STRING pathway analyses and functional enrichments identified genes involved in “mitochondrial part” (GO:0044429) with false discovery rate of 0.00094. **f)** KEGG pathway analyses identify genes enriched in PPARγ signaling pathway (false discovery rate 1.20e–05). **g)** Representative immunofluorescence of rat Schwann cell – mouse neuronal myelinated cocultures treated with 100 μM or 300 μM Arachidonic acid (complexed to BSA) for additional 7 days after 14 days in myelinating conditions. At the end of the treatment cultures were fixed and stained for MBP (rhodamine) and Neurofilament (fluorescein). Myelin degeneration was observed upon 300 μM Arachidonic acid treatment with no signs of axonal 0swelling. The same amount of BSA was added to control cocultures. Bar: 100 μm. **h, i**) qRT–PCR analyses on mRNA prepared from rat Schwann cell – mouse neuronal cocultures treated for additional 7 days with 100 μM or 300 μM Arachidonic acid after 14 days in myelinating conditions. Arachidonic acid induces the expression of *Pdk4* (**h**) and *Hmgcs2* (**i**) mRNA. Data have been normalized to *36B4* (*Pdk4*) and *TBP* (*Hmgcs2*) expression level and analyzed with the CFX Manager Software^TM^ from Biorad. N = 3 different mRNA preparations and analyses. Error bars represent mean ± s.e.m. (Unpaired t-test. *Pdk4*: **** P <0.0001, t=38.16, df =4; *Hmgcs2*: * P = 0,0429, t=2.92, df =4).

To validate these results, we performed quantitative RT–PCR analyses for some of the identified genes. We confirmed upregulated expression of *Pyruvate dehydrogenase kinase 4* (*Pdk4)* and of *Hydroxymethylglutaryl–CoA synthase* (*Hmgcs2)*, not only in myelinated cocultures treated with AT–56 (Fig. 5b), but also in sciatic nerves of *P0–Cre//l–pgds^flx/flx^*mice (Fig. 5c). Further, also *Acyl–CoA Synthetase Long Chain Family Member 3 (Acsl3)* and *Angiopoietin–like 4 (Angptl4)*, genes encoding for enzymes involved in lipids and Co–enzyme A consumption or release, are specifically upregulated in rat Schwann cells (Fig. 5b), indicating a possible rewiring in cell metabolism during myelin degeneration both *in vitro* and *in vivo*. Interestingly, STRING analysis revealed a strong inter–connection between proteins encoded by nine of these genes with a PPI enrichment p–value < 1.0e–16, suggesting that these proteins are at least partially biologically associated in their function. Moreover, analysis of the functional enrichments, designated genes involved in “cellular response to fatty acid” (GO:0071398) and in “regulation of lipid metabolic process” (GO:0019216) both with a false discovery rate of 2.13e–05 (Fig. 5d). Analysis of cellular component associated genes identified “mitochondrial part” (GO:0044429) with false discovery rate of 0.00094 (Fig. 5e). Finally, analysis of KEGG pathways revealed genes enriched in PPARγ signaling pathway with a false discovery rate of 1.20e–05 (Fig. 5f).

### Excess of Arachidonic acid is sufficient to induce myelin degeneration in vitro

Lysophosphatidylcholines derive from the release of one fatty acid chain from phosphatidylcholines and contribute to plasma membrane lipids recycling. They can in fact be further cleaved to release the other fatty acid chain from the choline group; alternatively, they can bind a free fatty acid to create a new phosphatidylcholine that will eventually relocate in the plasma membrane. Since in our *in vitro* and *in vivo* lipidomic analyses both lysophosphatidylcholine containing long fatty acid chain and phosphatidylcholine were decreased, we reasoned unlikely their new insertion into the plasma membrane. Rather, we speculated that the decrease in lysophosphatidylcholine might indicate an intracellular accumulation of free omega–6 fatty acids that could directly contribute to the morphological alterations observed in the absence of L–PGDS. To corroborate this hypothesis, we exploited our *in vitro* culture system and treated already myelinated mouse DRG–rat Schwann cell cocultures with Arachidonic acid, the omega 6 fatty acid direct precursor of prostanoid synthesis. We specifically added 100 μM or 300 μM Arachidonic acid bound to bovine serum albumin (BSA), as in this form, the Arachidonic acid crosses the plasma membrane and accumulates in the cytoplasm without exiting the cell. Although 100 μM Arachidonic acid–BSA is sufficient to slightly alter myelin integrity, addition of 300 μM Arachidonic acid–BSA causes extensive myelin degeneration, without signs of axonal swelling (Fig. 5g) or Schwann cell death (Supplementary Fig. 5b), as observed when L–PGDS enzymatic activity is inhibited^16^. These results strongly indicate that myelin degeneration is a direct consequence of Arachidonic acid accumulation at least *in vitro*. Further, in myelinated cocultures treated with Arachidonic acid we observed a specific upregulation in the mRNA of *Pdk4* (Fig. 5h) and *Hmgcs2* (Fig. 5i), two genes identified in the transcriptomic analyses, which are also upregulated in sciatic nerves of *P0–Cre//l–pgds^flx/flx^* mice.

Thus, the decrease in lysophosphatidylcholines and in phosphatidylcholines–containing long chain unsaturated fatty acid causes myelin degeneration in the absence of *l–pgds in vivo* in Schwann cells or *in vitro* when L–PGDS is not enzymatically active. Moreover, our *in vitro* data suggest that accumulation of Arachidonic acid, the substrate of L–PGDS, is sufficient to cause myelin alteration, by possibly altering Schwann cell metabolism.

### Loss of l–pgds enzymatic activity determines metabolic rewiring causing myelin degeneration

In our *in vivo* analyses we observed impairment in glycolysis as well as in the Krebs cycles starting at 6 months of age in *l-pgds^-/-^* mice. Interestingly, in *P0–Cre//l–pgds^flx/flx^* mice, acetyl–CoA levels started to diminish at 10 months (ctrl: 0.650 ± 0.15 vs mutant: 0.269 ± 0.077 P = 0.0171) and even more significantly at 14 months (ctrl: 0.508 ± 0.032 vs mutant: 0.176 ± 0.033 P = 0.0009), indicating an impairment of Krebs cycle in sciatic nerves of mice lacking glial L–PGDS expression.

To clarify the origins of these metabolic alterations, we performed metabolic flux analyses on Schwann cells – DRG neuronal myelinated cocultures in which we inhibited L–PGDS enzymatic activity as above described. 24 hours before cells’ harvesting, we added either 1 mM [U–^13^C_6_]–glucose, 2 mM [U–^13^C_5_]–glutamine, 200 μM [U–^13^C_16_]–Palmitate or 200 μM [U–^13^C_2_]–Sodium acetate to regular culture medium, then we followed labeled metabolites by LC–MS/MS analyses. To avoid any confounding results, we performed these experiments in physiological conditions, thus maintaining unaltered both glucose and glutamine final concentrations.

### Glucose metabolism

We did not detect significant differences in glycolysis flux (Fig. 6a); specifically, M+6 glucose, M+6 glucose–6P, M+6 fructose–bis–P isotopologues of the preparatory stage of glycolysis were comparable to controls, as well as pay–off phase isotopomers M+3 GAP/DHAP, M+3 phosphoenolpyruvate (PEP) and M+3 pyruvate. Of note, we observed a significant decrease in the acetyl–CoA (Fig. 6c) deriving from [U–^13^C_6_]–glucose that could be either due to excessive consumption or limited production. Since citrate M+2 and α–ketoglutarate M+2 levels were comparable in AT–56 and DMSO treated myelinated cocultures (Fig. 6e), the decrease in acetyl–CoA could be caused by a limited production rather than excessive consumption.

**Figure 6:**
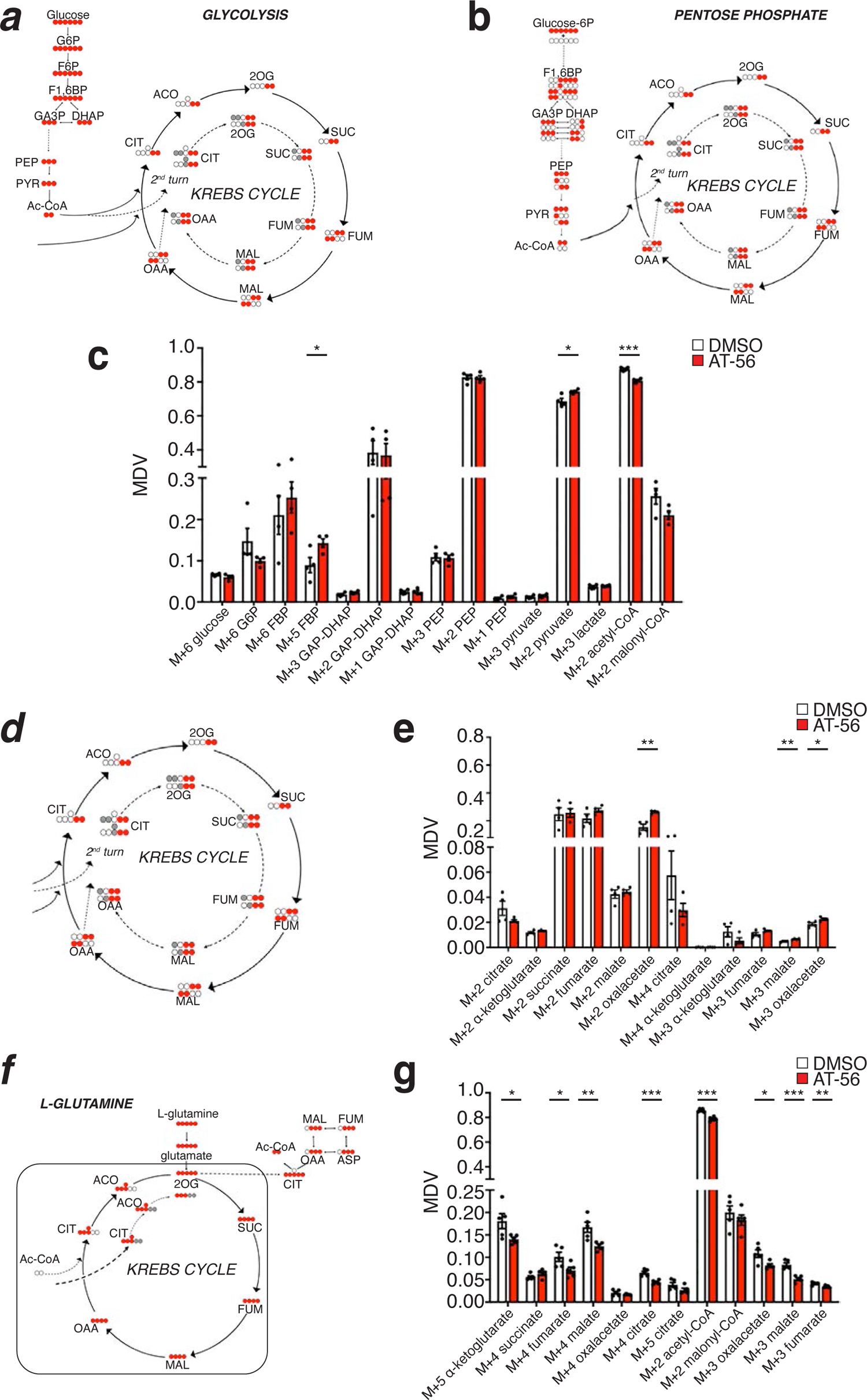
In vitro flux analyses: glucose and L–glutamine metabolism. Schwann cell – neuronal myelinated cocultures were allowed to myelinate for 14 days and then treated for additional 7 days with 25 μM AT–56 or with DMSO as vehicle control. Each labeled metabolite was added 24 hours before processing the samples in physiological conditions. The acetyl–CoA deriving from glucose and L–glutamine metabolism does not sustain the Krebs cycle in AT–56 treated cocultures. **a)** Metabolic fluxes scheme for [U–^13^C_6_]–glucose metabolism (glycolysis) showing the distribution of labeled carbons. (G6P: glucose–6–phosphate; FBP: fructose–6–phosphate; GAP: glyceraldehyde 3-phosphate DHAP: dihydroxyacetone phosphate PEP: phosphoenolpyruvate). **b)** Metabolic fluxes scheme for [U–^13^C_6_]–glucose metabolism (pentose phosphate) showing the distribution of labeled carbons. **c)** Graph showing the relative abundance of the labeled metabolites deriving from [U–^13^C_6_]–glucose, presented as a mass distribution vector (MDV). N = 4 different independent coculture experiments. Error bars represent mean ± s.e.m. (Unpaired t-test. M+5 FBP *P = 0.0444, t=2.534, df =6; M+2 pyruvate *P = 0.0198, t=3.15, df =6; M+2 acetyl–CoA ***P = 0.0002, t=8.218, df =6; all other metabolites tested not significantly different). **d)** Metabolic fluxes scheme for [U–^13^C_6_]–glucose metabolism (Krebs Cycle) showing the distribution of labeled carbons. **e)** Graph showing the relative abundance of the labeled metabolites deriving from [U–^13^C_6_]–glucose and entering the Krebs cycle, presented as a mass distribution vector (MDV). N = 4 different independent coculture experiments. Error bars represent mean ± s.e.m. (Unpaired t-test. M+2 oxalacetate **P = 0.0025, t=4.966, df =6; all other metabolites tested not significantly different). **f)** Metabolic fluxes scheme for [U–^13^C_5_]–glutamine metabolism showing the distribution of labeled carbons. **g)** Graph showing the relative abundance of the labeled metabolites deriving from [U–^13^C_5_]–glutamine presented as a mass distribution vector (MDV). N = 4 different independent coculture experiments. Error bars represent mean ± s.e.m. (Unpaired t-test. M+5 α–ketoglutarate *P = 0.0478, t=2.334, df =8; M+4 fumarate *P = 0.0403, t=2.445, df =8; M+4 malate **P = 0.0082, t=3.492, df =8; M+4 citrate ***P = 0.0006, t=5.424, df =8; M+2 acetyl–CoA ***P = 0.0001, t=7.112, df =8; M+3 oxalacetate *P = 0.0205, t=2.879, df =8; M+3 malate ***P = 0.0005, t=5.703, df =8; M+3 fumarate **P = 0.0057, t=3.743, df =8; all other metabolites tested not significantly different).

Analyses of different isotopomers of the same metabolites originating from the pentose phosphate pathway revealed an accumulation of M+2 pyruvate (Fig. 6b–c), which is consistent with the decrease in Acetyl CoA production (Fig. 6c). These results are strongly corroborated by a limited activity of PDH, which correlates with the observed transcriptomic increase in *Pdk4* expression. PDK4, in fact, inhibits PDH activity and limits acetyl–CoA production from pyruvate, thereby regulating metabolite flux through the Krebs cycle. Interestingly, despite the reduction in acetyl–CoA, all other metabolic intermediates of the Krebs cycle were not altered (Fig. 6d–e). Indeed, we observed an increase in M+2 oxaloacetate (Fig. 6e) that is synthetized from pyruvate through pyruvate carboxylase, an alternative metabolic strategy to replace oxaloacetate, occurring when PDH is blocked ^25^.

### L–Glutamine metabolism

Next, we followed the metabolic fate of [U–^13^C_5_]–glutamine in Schwann cells – neuronal myelinated cocultures treated with 25 μM AT–56 as compared to DMSO vehicle only treated controls. Glutamine is catabolized by either the mitochondrial oxidative pathway or by the cytosolic reductive branch. In AT– 56 treated cocultures, we observed a significant decrease in M+5 α–ketoglutarate, an isotopologue common to both glutamine catabolic pathways. Similarly, we found a reduction in M+4 fumarate, M+4 malate and M+4 citrate isotopologues of the oxidative glutamine pathways (Fig. 6f–g). These data indicate limited glutamine oxidative catabolism in the Krebs cycle. Further, we observed an overall decrease activity of the glutamine reductive pathway, as M+3 fumarate, M+3 malate and M+3 oxalacetate levels were also reduced, while M+5 citrate was unaffected in the absence of L–PGDS (Fig. 6g). Of note, M+2 acetyl–CoA levels, resulting from both glutamine catabolic pathways, were reduced (Fig. 6g). Thus, blocking L–PDGS enzymatic activity impairs both glutamine oxidative and reductive metabolism.

### Fatty acid β–oxidation

The decrease in acetyl–CoA production deriving from glucose and glutamine metabolism, coupled to *in vivo* lipids’ dysregulation and *Acsl3* and *Cpt1a* upregulation observed in transcriptomic analyses, would suggest an increase in fatty acids β–oxidation. Acsl3, in fact, binds specifically free long–chain fatty acids to CoA determining either their degradation or incorporation in cellular lipids. On the contrary, Cpt1a catalyzes the transfer of the acyl group of long–chain fatty acid–CoA conjugates onto carnitine, an essential step for mitochondrial uptake of long–chain fatty acids and their β–oxidation. Thus, to determine whether in the absence of functional L–PGDS β–oxidation is hampered, we followed its metabolic flux by treating degenerating cocultures with 200 μM [U–^13^C_16_]–palmitate (Fig. 7a).

**Figure 7:**
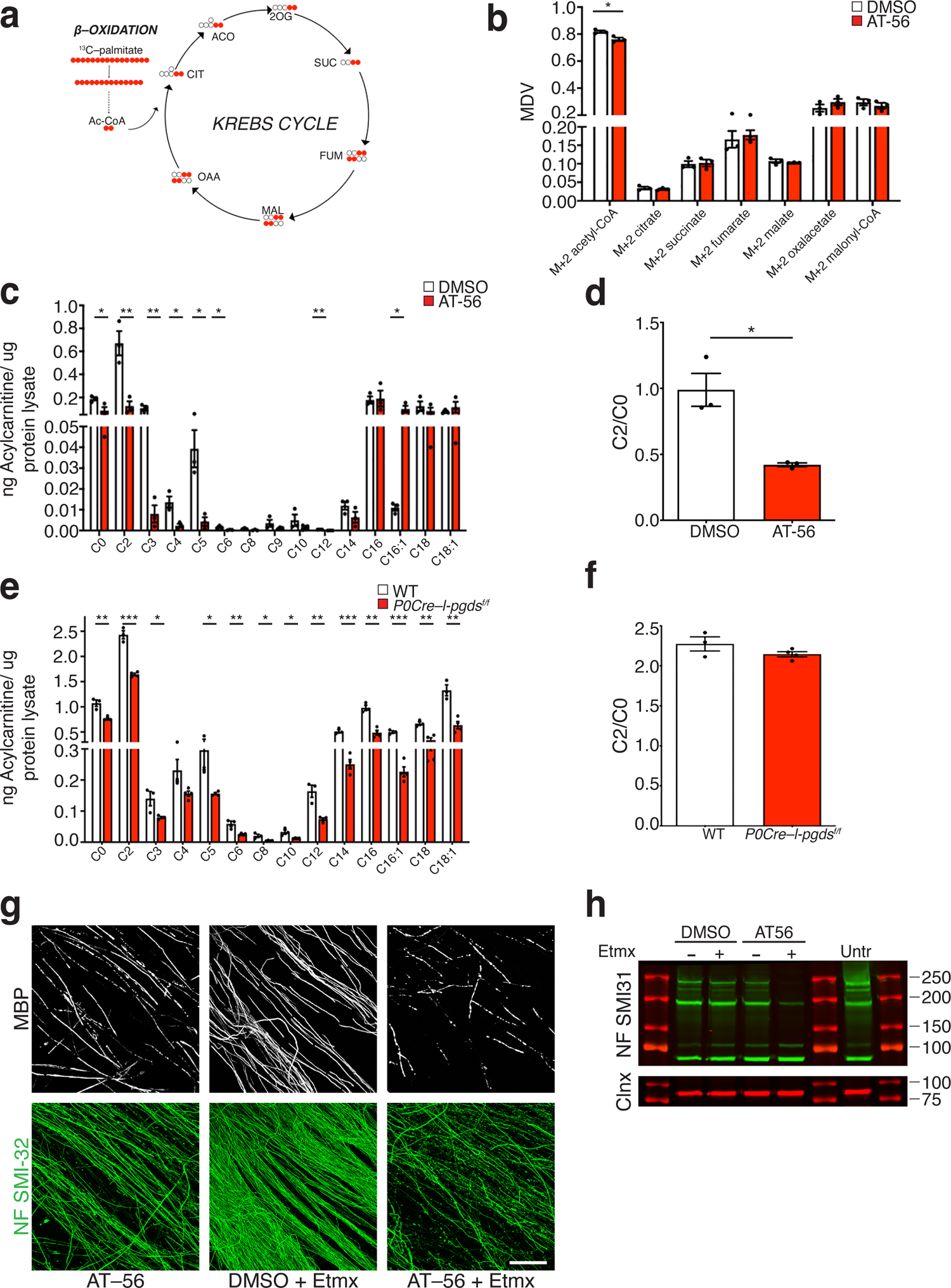
Fatty acids β–oxidation in the absence of L–PGDS. **a)** Metabolic fluxes scheme for [U–^13^C_16_]–palmitate metabolism showing the distribution of labeled carbons. b) Graph showing the relative abundance of the labeled metabolites deriving from [U–^13^C_16_]–palmitate presented as a mass distribution vector (MDV) in Schwann cell – neuronal cocultures myelinated for 14 days and then treated for additional 7 days with 25 μM AT–56 or with DMSO as vehicle control. Labeled palmitate was added 24 hours before processing the samples in physiological conditions. The acetyl– CoA deriving from β–oxidation metabolism does not sustain the Krebs cycle in AT–56 treated cocultures. N = 3 different independent coculture experiments. Error bars represent mean ± s.e.m. (Unpaired t-test. M+2 Acetyl–CoA *P = 0.0225, t=3.614, df =4; all other metabolites were not significantly different). c) Graph showing levels of acyl–carnitines in Schwann cell – neuronal myelinated cocultures treated with 25 μM AT–56, normalized over the total protein content. N = 3 different independent coculture experiments. Error bars represent mean ± s.e.m. (Unpaired t-test. C0 *P = 0.0413, t=2.967, df =4; C2: **P = 0.0085, t=4.824, df =4; C3: **P = 0.0082, t=4.877, df =4; C4: *P = 0.0168, t=3.952, df =4; C5: *P = 0.0189, t=3.814, df =4; C6: *P = 0.022, t=3.638, df =4; C12: **P = 0.0011, t=8.485, df =4; C16:1: *P = 0.0313, t=3.254, df =4). d) Graph showing the ratio between C2/C0 acyl–carnitines, a hallmark of fatty acid β oxidation, in Schwann cell – neuronal myelinated cocultures treated with 25 μM AT–56 or with DMSO as vehicle control. N = 3 different independent coculture experiments. β oxidation is reduced when L–PGDS is inactive. Error bars represent mean ± s.e.m. (Unpaired t-test. *P = 0.0107, t=4.518, df =4). e) Graph showing levels of acyl–carnitines in 14 months old wild type littermate controls (WT) and *P0– Cre//l–pgds^flx/flx^* sciatic nerves, normalized over the total protein content. N = 4 *P0–Cre//l–pgds^flx/flx^* mice;/ N = 3 littermate controls (WT) mice. Error bars represent mean ± s.e.m. (Unpaired t-test. C0 **P = 0.003, t=5.357, df =5; C2: ***P = 0.0001, t=10.8, df =5; C3: *P = 0.0267, t=3.106, df =5; C5: *P = 0.0466, t=2.628, df =5; C6: **P = 0.0062, t=4.535, df =5; C8: *P = 0.0149, t=3.639, df =5; C10: *P = 0.0168, t=3.529, df =5; C12: **P = 0.0032, t=5.298, df =5; C14: ***P = 0.0005, t=8.064, df =5; C16: **P = 0.001, t=6.81, df =5; C16:1: ***P = 0.0002, t=9.699, df =5; C18: **P = 0.0031, t=5.327, df =5; C18:1 **P = 0.0026, t=5.576, df =5). f) Graph showing the ratio between C2/C0 acyl–carnitines, a hallmark of fatty acid β oxidation, in 14 months old wild type littermate controls (WT) and *P0–Cre//l–pgds^flx/flx^*sciatic nerves. N = 4 *P0–Cre//l– pgds^flx/flx^* mice; / N = 3 littermate controls (WT) mice. β oxidation does not change in the absence of L-PGDS. Error bars represent mean ± s.e.m. (Unpaired t-test. P = 0.1824 n.s., t=1.548, df =5). g) Representative immunofluorescence of 14 days myelinated Schwann cell – neuronal cocultures treated for additional 9 days with 25 μM AT–56, in the presence or absence of 50 μM etomoxir, a specific *Cpt1a* inhibitor. At the end of the treatment cultures were fixed and stained for MBP (white) and non-phosphorilated neurofilament, SMI–32 (fluorescein). Unlike vehicle only treated cocultures (DMSO + 50 μM etomoxir), blocking *L–pgds* and *Cpt1a* causes myelin and axonal degeneration. Bar: 100 μm. h) Representative western blotting analyses of 14 days myelinated Schwann cell – neuronal cocultures treated for additional 9 days with 25 μM AT–56, in the presence or absence of 50 μM etomoxir, a specific *Cpt1a* inhibitor. Lysates were tested for phosphorylated neurofilament (SMI–31) and Calnexin (Clnx) as a loading control. Blocking *L–pgds* and *Cpt1a* significantly impairs neurofilament expression.

Levels of acetyl–CoA deriving from palmitate β–oxidation were reduced (Fig. 7b), suggesting that this metabolite could be either more consumed or less produced from fatty acids catabolism. Also in this case, we did not observe increased amounts in any of the Krebs cycle metabolites deriving from labeled palmitate (Fig 7b), making unlikely that the reduction in acetyl–CoA is due to excessive consumption in this pathway. To corroborate these results, we quantified the acylcarnitine pool *in vitro* in Schwann cell– neuronal cocultures treated with AT–56 (Fig. 7c) and in vivo in *P0–Cre//l–pgds^flx/flx^*sciatic nerves’ lysates (Fig. 7e). However, in both conditions, the ratio of acetyl-carnitine (C2)/free carnitine (C0), a hallmark of overall fatty acid β-oxidation, was unchanged or even decreased (Fig. 7d, 7f).

Palmitate tracing experiment indicates that acetyl-CoA levels were reduced in cocultures in the absence of functional L–PGDS (Fig. 7b). To understand the importance of fatty acid derived acetyl– CoA, we inhibited fatty acid entrance into mitochondria by blunting Cpt1a enzymatic activity, and hence reducing mitochondrial acetyl–CoA production, in Schwann cells neuronal cocultures treated with AT–56. Addition of 50 μM Etomoxir, a specific Cpt1a inhibitor ^26^, together with 25 μM AT–56 to already myelinated cocultures, strikingly compromised axonal integrity as shown by immunohistochemical analyses for MBP and non-phosphorylated neurofilament (SMI32) (Fig. 7g). Notably, in these conditions, the expression level of phosphorylated neurofilament (SMI31) was also significantly hampered (Fig. 7h). Collectively, these results indicate that in the absence of a catalytically active L–PGDS, fatty acids– derived acetyl–CoA is required by Schwann cells to support axonal integrity.

### Krebs cycle is sustained by ketone bodies and acetate utilization in the absence of l–pgds

The acetyl–CoA deriving from fatty acid β–oxidation, which does not directly replenish the Krebs cycle (Fig. 7a, b) is probably used in other mitochondrial metabolic pathways such as ketone bodies biosynthesis. All of this is supported by the upregulation in the expression of *Hmgcs2* we observed *in vitro* and *in vivo* in *P0–Cre//l–pgds^flx/flx^* sciatic nerves (Fig. 5b–c). This enzyme catalyzes the condensation of acetyl–CoA into acetoacetyl–CoA to form HMG–CoA, which, in mitochondria, drives β–hydroxybutyrate synthesis, a ketone body carrier of energy to peripheral tissues in conditions of energy deprivation ^27, 28^. To assess whether in the absence of functional L–PGDS, Schwann cells and neurons might rely on ketone bodies as energy source, we measured β–hydroxybutyrate and acetoacetate levels *in vivo* in *P0–Cre//l–pgds^flx/flx^* sciatic nerves and in AT–56 treated cocultures. Interestingly, levels of β– hydroxybutyrate were significantly upregulated in 11 months *P0–Cre//l–pgds^flx/flx^*sciatic nerves (Fig. 8a). Similarly, we observed enhanced a release of β–hydroxybutyrate in conditioned media of AT–56 treated degenerated cocultures (Fig. 8b). Thus, it is possible that the release of β–hydroxybutyrate might accumulate in sciatic nerves to sustain energy production in the absence of *l–pgds*. Of note, we also observed a significant decrease in intracellular acetoacetate (Fig. 8c) along with reduced levels of succinyl–CoA, a key CoA donor in the ketone bodies utilization pathway (Fig. 8d), suggesting that acetoacetate produces acetyl–CoA likely re-entering the Krebs cycle, at least *in vitro*. In agreement, we found no alterations in ATP energy charge in *P0–Cre//l–pgds^flx/flx^* sciatic nerves (Fig. 8e) and in AT–56 treated cocultures (Fig. 8f). These results, along with the upregulation of *Acsl3* and *Cpt1a* expression and activity, strongly hint at a high rate of fatty acid β–oxidation of myelin lipids to sustain ketone bodies biosynthesis and their utilization as energy source. Moreover, since the acetyl–CoA deriving from glucose and L–glutamine does not sustain the Krebs cycle (Fig 6), we hypothesized that the necessary acetyl–CoA might derive from other sources. To determine whether the acetate metabolism (Fig. 8g), an important metabolic route of the nervous system when glucose is not available ^29^, could contribute to Krebs cycle functioning, we supplemented Schwann cell neuronal cocultures treated with AT–56 for 7 days, with 200 uM [U–^13^C_2_]–acetate using the same experimental paradigm above described. Although we detected a decrease in M+2 acetyl–CoA, we surprisingly observed a strong increase in M+3 citrate, M+2 and M+3 α–ketoglutarate, M+3 fumarate, and M+2 and M+3 oxaloacetate (Fig. 8h) deriving from labeled acetate, suggesting a more pronounced acetate consumption, which ensures, at least, Schwann cells and neuronal survival.

**Figure 8:**
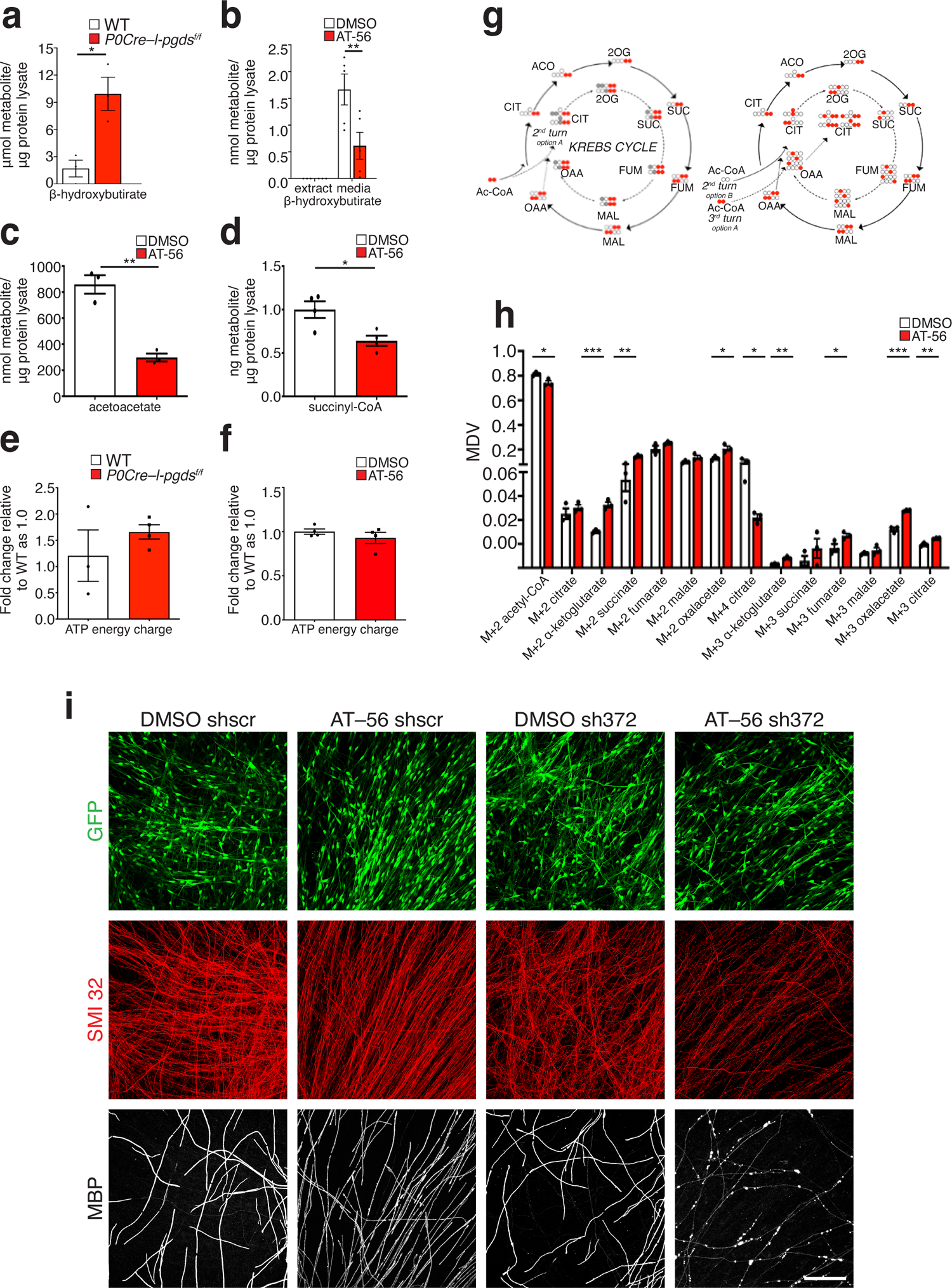
In the absence of L–PGDS myelinating Schwann cells increase ketone bodies synthesis and consumption. **a)** Graph showing that β–Hydroxybutyrate levels are significantly increased in lysates of 11 months old *P0–Cre//l–pgds^flx/flx^* sciatic nerves as compared to wild type littermate controls (WT). N = 3 different preparations. Error bars represent mean ± s.e.m. (Unpaired t test *P = 0.0157, t=4.029, df =4). **b)** Graph showing that β–Hydroxybutyrate levels are significantly reduced in the supernatant of Schwann cell – neuronal myelinated cocultures treated with 25 μM AT–56 as compared to DMSO vehicle controls, but not detectable in corresponding cell lysates. N = 5 from 2 different independent coculture experiments (Cell media) N = 3 cell lysates. Error bars represent mean ± s.e.m. (Two-way Anova. **P = 0.0077, F(4,4) = 18.44, df =4). **c)** Graph showing that acetoacetate levels are significantly reduced in Schwann cell – neuronal myelinated cocultures treated with 25 μM AT–56 as compared to DMSO vehicle controls. N = 3 different independent coculture experiments. Error bars represent mean ± s.e.m. (Unpaired t-test. **P = 0.0019, t=3.748, df =4). **d)** Graph showing that succinyl–CoA levels are significantly reduced in Schwann cell – neuronal myelinated cocultures treated with 25 μM AT–56 as compared to DMSO vehicle controls. N = 3 different independent coculture experiments. Error bars represent mean ± s.e.m. (Unpaired t-test. *P = 0.0189, t=3.189, df =4). **e).** Graph showing that the energy balance is similar in 14 months old *P0–Cre//l–pgds^flx/flx^* sciatic nerves as compared to wild type littermate controls (WT). N = 4 *P0–Cre//l–pgds^flx/flx^*mice; / N = 3 littermate controls (WT) mice. Error bars represent mean ± s.e.m. **f).** Graph showing that the energy balance is comparable in Schwann cell – neuronal myelinated cocultures treated with 25 μM AT–56 or with DMSO, as vehicle control. N = 4 different independent coculture experiments. Error bars represent mean ± s.e.m. g) Metabolic fluxes scheme for [U–^13^C_2_]–sodium acetate metabolism showing the distribution of labeled carbons. h) Graph showing the relative abundance of the labeled metabolites deriving from [U–^13^C_2_]–acetate presented as a mass distribution vector (MDV) in Schwann cell – neuronal cocultures myelinated for 14 days treated for additional 7 days with 25 μM AT–56 or with DMSO as vehicle control. Labeled acetate was added 24 hours before processing the samples in physiological conditions. N = 3 different independent coculture experiments. Error bars represent mean ± s.e.m. (Unpaired t-test. M+2 acetyl– CoA *P = 0.02, t=3.748, df =4; M+2 α–ketoglutarate ***P = 0.0009, t=8.953, df =4; M+2 succinate **P = 0.0034, t=6.217, df =4; M+2 oxalacetate *P = 0.0216, t=3.66, df =4; M+4 citrate *P = 0.0215, t=3.666, df =4; M+3 α–ketoglutarate **P = 0.0027, t=6.628, df =4; M+3 fumarate *P = 0.0496, t=2.784, df =4; M+3 oxalacetate ***P = 0.001, t=8.688, df =4; M+3 citrate **P = 0.0048, t=5.657, df =4; all other metabolites tested not significantly different). i) Representative immunofluorescence of wild–type myelinated Schwann cells neuronal cocultures infected with *Hmgcs2 sh372* or scramble shRNA (*shscr*). Three days post infection cocultures were supplemented with ascorbic acid for 10 days to allow myelination and then treated with 25 uM AT–56 or vehicle only, as control, for additional 7 days. At the end of the treatment, cultures were fixed and stained for GFP to visualize infected cells (fluorescein), non–phosphorylated neurofilament SMI–32 (rhodamine), and MBP (far-red). Ablation of *Hmgcs2* coupled to L–PGDS block, further hampers myelin integrity and determines axonal instability. *N* = 3 independent biologic replicates. Bar: 100 μm.

As above mentioned, one of the key enzymes for ketone bodies formation, is Hmgcs2, whose expression is upregulated *in vitro* and *in vivo* in the absence of L–PGDS (Fig. 5). One of the most striking results of this and our former study ^16^ is the morphological alteration in myelin not accompanied by axonal swallowing and/or neuronal cell death. To confirm that, in the absence of L–PGDS, Schwann cells activate an alternative pathway relying on ketone bodies production to sustain the axo-glial unit integrity, we ablated *Hmgcs2* expression by lentiviral mediated knocked down. Established Schwann cells–DRG neuronal cocultures, were infected with three different shRNA specifically designed against *Hmgcs2*. Three days post infection, cocultures were supplemented with ascorbic acid for additional 10 days to allow myelination and then treated with 25 uM AT–56 or vehicle only, as control, for additional 7 days. At the end of the treatment, cultures were fixed and stained for MBP, non-phosphorylated neurofilament and GFP to visualize infection’s levels. As expected, most cells infected by the shRNAs were Schwann cells (Fig. 8i). Importantly, knock down of *Hmgcs2*, which was validated by qRT–PCR analyses (Supplementary Fig. 6a), not only enhances myelin breakdown, but also dramatically hampers axonal integrity (Fig. 8i and Supplementary Fig. 6b).

Collectively, we posit that in the absence of glial L–PGDS, myelin lipid composition is altered, and with time, glial cells undergo a rewiring in metabolism. Indeed, while in early phases of myelin degeneration Schwann cells rely on glucose to sustain their energy balance, when degeneration continues Schwann cells switch to ketone bodies and acetate utilization to sustain the Krebs cycle and uphold the axo–glial unit (Supplementary Fig. 7).

## DISCUSSION

Metabolic homeostasis is center to the maintenance of a healthy and functional organ. This is particularly relevant in the case of the nervous system, where the cross communication between glial cells and neurons is essential to integrate a continuous transmission of electric impulses from the center to the periphery and vice versa ^1, 23^.

To thoroughly investigate the complex regulation of metabolism in nerves, we combined multiple approaches. We described and analyzed metabolites and lipids *in vivo* in sciatic nerves of three mutant mice at different age: a complete null L-PGDS, a Schwann cell specific and a motor neuron specific mutant. Next, we performed a systematic analysis of different metabolic pathways *in vitro* using the Schwann cell neuronal coculture system. Though *in vitro* we could partly reproduce the complexity of the interactions occurring in nerves, where in addition to glial cells and axons also other cells can contribute to metabolic changes, these studies were instrumental to identify key changes in metabolism that we confirmed in sciatic nerves.

We also report that L–PGDS expressed in Schwann cells or neurons exert different roles, with the neuronal enzyme regulating the thickness of the myelin sheath, in agreement with the model proposed in^16^. In turn, glial L-PGDS is key in controlling myelin homeostasis. Indeed, our results indicate that this enzyme in Schwann cells is crucial to maintain myelin lipid integrity, as, in *P0–Cre//l–pgds^flx/flx^* mice we observed a drastic change in myelin lipid composition. In particular, lysophosphatidylcholines, containing omega–6 fatty acids, which are at the basis of the prostaglandin synthesis, are released from myelin and likely accumulate in Schwann cells cytoplasm. Accordingly, *in vitro* increased levels of Arachidonic acid are sufficient to determine myelin degeneration and to dysregulate the expression of Pdk4 and Hgmcs2, key enzymes in controlling cellular energy metabolism.

Notably, changes in myelin lipids’ composition occur primarily in older mice and correlate with accumulation of morphological changes. This effect is particularly compelling in *P0–Cre//l–pgds^flx/flx^* sciatic nerves in which they are mostly evident. Thus, we propose that L–PGDS might be functional either to maintain intact the myelin structure or be part of a coordinated mechanism controlling myelin homeostasis and remodeling. Since, we did not observe any significant overt morphological phenotype in mice before 6–months of age in *l–pgds^-/-^* mice and even later in *P0–Cre//l–pgds^flx/flx^* sciatic nerves, we favor the last assumption, although we cannot exclude that lipids’ remodeling could be more efficient in younger animals. Indeed, lipids and metabolites dysfunctions precede myelin alterations both in 4 months old *l–pgds^-/-^* mice and in 8 moths old *P0–Cre//l–pgds^flx/flx^* mice.

Based on the results of our study, we envisage two different hypotheses to explain how the absence of L–PGDS alters myelin morphology. Although in adult mice myelin lipids’ remodeling could be less efficient, it is also possible that the demand of PGD2 in Schwann cells induces a continuous release of lipids from the plasma membrane, reaching a level above which the structural integrity of myelin cannot be maintained. Regardless of the mechanisms, we have identified the synthase L–PGDS as central in maintaining an undamaged myelin. In future studies it might be important to define whether L–PGDS cooperates with other molecules in this task, and whether they could be pharmacologically targeted to minimize myelin alterations. Interestingly, dietary supplementation of phosphatidylcholine and phosphatidylethanolamine, the main class of lipids we observed reduced in our studies, improved myelination and nerve conduction velocity in a rat model of CMT1A ^30^.

A remaining open question is the final destiny of the lipids released from the plasma membrane. Accumulation of arachidonic acid in Schwann cells is sufficient to upregulate *pdk4* and *Hmgcs2* expression, whose expression are also upregulated in *P0–Cre//l–pgds^flx/flx^* sciatic nerves. Upregulation of *pdk4* might therefore be instrumental to limit acetyl–CoA production by inactivating the pyruvate dehydrogenase, slowing down Schwann cells glycolysis. Indeed, we observed a general attenuation in the use of glucose as the main source to preserve the axo–glial unit both *in vitro* and *in vivo*. In this context, it is important to note that previous studies have shown that the acetyl–CoA deriving from pyruvate is dispensable for myelin maintenance and axon integrity ^31^.

Surprisingly, in *l–pgds^-/-^* mice we also found a less efficient lactate production and Schwann cells rely more on acetate. Though lactate is generally transferred to axons to sustain axonal metabolic demand^32^, its role in sustaining the energetic metabolism of peripheral nerves is complex and controversial ^33, 34^.

It is thus tempting to speculate that acetate might be a specific metabolite used to support neurons and glial cells as a substitute for lactate in adulthood ^34^. Whether this switch in metabolites’ production occurs only in adulthood or it is present also in development and/or in pathological settings requires additional investigations.

Our results strongly indicate that continuous remodeling of plasma membrane myelin lipids, due to the absence of L–PGDS, drives Schwann cells towards a change in their metabolic activity. The decreased mitochondrial fatty acid–derived acetyl–CoA, is likely used to generate ketone bodies to boost ketolysis ^18^, which serves as an alternative energetic route for lipid synthesis, especially in the nervous system ^18, 35^. In both CNS ^36^ and PNS ^37^ ketone bodies are preferentially used for myelin lipid synthesis, mainly cholesterol. Although, we did not observe any increase in malonyl–CoA consumption in the absence of L–PGDS in flux analyses, we cannot exclude that acetate might foster new lipid synthesis *in vivo* or might be preferentially activated in young mutant mice. Of note, metabolic incorporation of acetate into PNS myelin is prevalently used very early in development and after injury ^38^, and in Trembler mutant mice, which are characterized by a hypomyelinating peripheral neuropathy, acetate represents the main source for lipid synthesis ^37^. It is therefore possible that ketolysis and acetate might be used also in our model to sustain myelin remodeling.

Since lipids can function as second messenger in the cells ^39, 40^, it is also possible that those released from the plasma membrane in the absence of L–PGDS could act onto PPARγ to promote gene transcription. Indeed, many of the genes we found upregulated in our transcriptomic analyses, are implicated in the regulation of lipid metabolic process. Finally, we cannot exclude that both systems co– exist. Hence, released lipids might on one side promote gene transcription to counteract continuous PNS myelin remodeling, and in parallel limit excessive production of acetyl–CoA in the mitochondria favoring ketolysis. In turn, in young mice, ketolysis could foster new lipid synthesis thus preventing excessive myelin remodeling.

It should be noted that the metabolic shift occurs essentially in glial cells, as *in vivo* analyses confirmed that *P0–Cre//l–pgds^flx/flx^*but not *Chat–Cre//l–pgds^flx/flx^* sciatic nerves presented aberrant myelin as well as metabolic changes overlapping those observed in *l–pgds^-/-^*mice. Though metabolic defects in *P0–Cre//l–pgds^flx/flx^* and *l–pgds^-/-^* mice are remarkably similar, they also present differences. In *P0–Cre//l–pgds^flx/flx^* nerves lipid alterations are significantly evident at 14 months, while they are already apparent at 6 months in *l–pgds^-/-^* mice. At metabolic level, both mutants present a striking reduction in glycolysis, with an abrupt acetyl–CoA reduction in *P0–Cre//l–pgds^flx/flx^* sciatic nerves. These differences are not surprising as L–PGDS is expressed in several cells in addition to Schwann cells and neurons. Alternatively, it is possible that the hypomyelinating phenotype present in *l–pgds^-/-^* mice but not in *P0–Cre//l–pgds^flx/flx^* sciatic nerves, might render myelin more susceptible to changes in lipid composition, thus enhancing the overall process. Yet, it is remarkable that ablation of glial L–PGDS is sufficient to trigger the metabolic shift in peripheral nerves. This observation is further corroborated by the *in vitro* results, in which, in a simplified system containing mostly neurons and Schwann cells, blocking L–PGDS enzymatic activity essentially reproduces results observed in *P0–Cre//l–pgds^flx/flx^* nerves. Indeed, we observed a selective upregulation of genes involved in fatty acids regulation exclusively in Schwann cells. On this basis, metabolic rewiring occurs in Schwann cells, especially the one attributable to ketogenesis fueled by fatty acids derived from degenerating myelin. Thus, we propose that the observed metabolic changes represent an adaptive response to ongoing morphological alterations that are ensuing in Schwann cells because of continuous myelin remodeling.

Our study also reveals that in the absence of L–PGDS Schwann cells rely on acetate to sustain the Krebs cycle and that levels of acetoacetate, succinyl–CoA and β–hydroxybutyrate are reduced in the absence of L–PGDS, further supporting the notion that Schwann cells undergo enhanced ketolysis when L–PGDS is not expressed or inactive. Notably, levels of β–hydroxybutyrate were diminished in cell culture media, but upregulated in *P0–Cre//l–pgds^flx/flx^* sciatic nerves (Fig. 8), suggesting that the release of β–hydroxybutyrate could accumulate in the nerve where it is used to produce ketone bodies and fuel the Krebs cycle in addition of indicating that the inter–conversion between acetoacetate and β– hydroxybutyrate is shifted towards the latter. Although we do not have formal proof for the transfer of β–hydroxybutyrate from Schwann cells to neurons to sustain their energy demand and the integrity of the axo–glial unit (Supplementary Fig. 7), we have several evidence supporting this hypothesis: i) in the absence of a functional L–PGDS we did not observe any impairment in neuronal survival ^16^(and this study); ii) ablation of *Hmgcs2*, a key enzyme for ketone bodies synthesis, as well as iii) block of lipid transfer into mitochondria, not only worsens myelin degeneration, but drastically compromises axonal stability. All these observations strongly support our hypothesis that ketone bodies synthesis in Schwann cells in the absence of L–PGDS, is critical to sustain the axo-glial unit. Interestingly, ketogenic diet reduces metabolic induced–allodynia and promotes nerve growth in the epidermis ^41^. Further, it improves CNS myelination and increases the number of oligodendrocytes in a model of Pelizaues–Merzbacher disease ^42^.

In conclusion, we have unveiled a new pathway connecting prostaglandin activity to Schwann cell metabolism and myelin homeostasis in adulthood. Based on the results of our studies, we propose that glial L–PGDS is part of a coordinated program aiming at preserving myelin integrity. Importantly, our study suggests that Schwann cell myelin lipids, particularly fatty acids, are a valuable reservoir to produce ketone bodies, which together with acetate represent the adaptive substrates they can rely on under threatening circumstances to sustain the axo-glial unit and preserve the integrity of the PNS.

## MATERIALS AND METHODS

### Mice and genotyping

All experiments were performed on male mutants and wild type mice in a C57/Bl6 congenic background following protocols approved by the Institutional Animal Care and Use Committee of San Raffaele Hospital and by the Italian Minister of Health (Protocol number 973). Generation of *l–pgds^-/-^, P0–CRE*, *Chat–CRE* mice and genotypes’ determination were previously described ^43^ ^20, 21, 44^. *l–pgds^flx/flx^* mice were genotyped by PCR analysis on genomic DNA, using the following primers: 5’–GGG CAC TGT CAG CCT GTG TGC TTG TGC –3’ (forward), 5’–CCA CAC AGG TCC TAG CAG CAT GCCTC–3’ (reverse). Cycling conditions were: 94°C for 60 s, 60°C for 60 s, and 72°C for 60 s (29 cycles), followed by a 10 min extension at 72°C.

For *Chat–CRE//l–pgds^flx/flx^* mice, the presence of the null allele was determined by PCR on genomic DNA extracted from ventral and dorsal spinal cord regions using the following primers: 5’–GGG CAC TGT CAG CCT GTG TGC TTG TGC–3’ (forward LoxP1), 5’– GGT GAG AGA AGT CAG TCA GAG GGC TGG–3’ (forward Lox P2) and 5’– CCT GGC TCC TTG GAG ACC CCT GCT GC–3’ (reverse Lox P2). Cycling conditions were: 94°C for 60 s, 61°C for 60 s and 72°C for 60 s (30 cycles), followed by a 3 min extension at 72°C. Amplified fragments were analyzed on a 2% agarose gel.

### Morphological and morphometric analyses

Semi–thin and ultrathin sections were obtained as described ^45^. Sciatic nerves were removed and fixed with 2% glutaraldehyde (Electron Microscopy Science) in 0.12 M phosphate buffer, post fixed with 1% osmium tetroxide (Electron Microscopy Science), and embedded in Epon (Epoxy Embedding Medium kit, Sigma). Semi–thin transversal sections (0.5–1 μm thick) were stained with toluidine blue and examined by light microscopy (Olympus BX51). Digitized non–overlapping images from corresponding levels of the sciatic nerve were obtained with a digital camera (Leica DFC300F) using a 100X objective. Images were merged and transversal section analyzed using the NIH ImageJ v1.45s software (NIMH Image Library, RRID:SCR_005588). The percentage of myelin aberrations was determined as the number of fibers presenting myelin structural alterations over the total number of fiber in the analyzed section. *g*–ratio measurements were performed on non–overlapping digitalized electron micrographs images as reported in ^46^ or using the Leica QWIN V3 software.

### Cell cultures and treatments

Mouse dorsal root ganglia (DRG) neurons were isolated from E14.5 embryos and established on rat collagen I (Cultrex) coated glass coverslips or 6–well plates as previously described ^17^. Cells were grown in Neuronal Basal media (Gibco), B27 (Gibco), 4g/L D–glucose (Fluka), 2 mM L–glutamine (Invitrogen), supplemented with 50 ng/ml NGF (B.5017, Harlan Laboratories, Indianapolis, IN). In some experiments explants were cycled with FUDR to eliminate all non–neuronal cells. Primary rat Schwann cells were prepared as described in ^17^ and maintained in DMEM (Invitrogen), 10% FBS (vol/vol, Invitrogen), 2 mM L–glutamine (Invitrogen), 2 μM Forskolin (Sigma) and 10 ng/ml rhNRG1 (R&D), until used. Rat Schwann cells were added (200.000 cells per coverslip) to established explant cultures of DRG neurons and cultured in MEM (Invitrogen), 10% FBS (vol/vol, Invitrogen), 2 mM L–glutamine (Invitrogen), 4 g/L D–glucose (Fluka) and NGF. Myelination was initiated by supplementing media with 50 μg/ml ascorbic acid (Sigma–Aldrich) and continued for 10–14 days.

Myelin degeneration was obtained by treating myelinated Schwann cell–DRG neuronal co-cultures with 25 μM AT–56 (Tocris) in myelinating medium for additional 7 days, unless otherwise specified. In Arachidonic acid experiments, myelinated rat Schwann cell–mouse DRG neuronal co-cultures were treated with different concentrations of Arachidonic acid (Cayman Chemical Company) for 7 days. Briefly, ethanol was evaporated under nitrogen flux and Arachidonic acid dissolved in DMSO to a final concentration of 164 mM. The fatty acid was bound to BSA with a molar ratio 1:5 by heating the stock solution 7% BSA / 5 mM Arachidonic acid at 37°C for 1 hour. The compound was then diluted in myelinating medium.

In experiments in which we blocked *Cpt1a* activity, we treated already myelinated Schwann cell– DRG neuronal cocultures with 25 μM AT–56 and 50 μM Etomoxir (Sigma–Aldrich) for 9 days. Cultures were then processed for immunofluorescence analyses or Western Blotting analyses as detailed below.

In metabolomics flux analyses, labeled metabolites were added to degenerating co–cultures 24 hours before collecting the cells. In details, L–glutamine in the culture medium was completely substituted with 2 mM [U–^13^C_5_]–glutamine, 1 mM [U–^13^C_6_]–glucose was added in medium containing 21.2 mM glucose (final total glucose 22.2 mM), [U–^13^C_16_]–Palmitate was added to 200 μM final concentration (together with 1 mM L-carnitine) and [U–^13^C_2_]–Sodium acetate to 2 mM.

### Lentiviral production and infection

Individual shRNA clones (FE5V3SM11241-231157720: sh372; FE5V3SM11242-240857410: sh062; FE5V3SM11242-243698710: sh362) specifically targeting mouse *Hmgcs2* were obtained from Dharmacon (Horizon Discovery). Lentiviral vectors were transfected into HEK293T cells (Dharmacon) and viruses were produced according to the Dharmacon Trans-Lentiviral Packaging System’s instructions. Briefly, cells confluent at 80% were transfected using Ca_2_P0_4_ and viral particles were collected 3 days post transfection by centrifugation at the ultracentrifuge at 4°C, 20.000 rpm for 2 hours (Beckman). Collected viral particles were then suspended in DMEM and stored to –80°C, until used. To assess viruses’ titer HEK293T cells were infected with different dilutions of each lentiviral vectors for 4 h and the number of turboGFP positive colonies was counted three days post infection. The viral titer was calculated as following: n colonies x dilution factor x 40 = TU/mL (transducing unit/mL) (Dharmacon’s instructions). Established Schwann cells–DRG neuronal cocultures one week post dissection were infected with lentiviruses at an MOI ranging between 5 and 9 and incubated for 16 h in MEM (Invitrogen), 5% FBS (vol/vol, Invitrogen), 4 mM L–glutamine (Invitrogen), 2 g/L D–glucose (Fluka) and 5 ng/mL NGF. Three days after infection we induced myelination by treating the cocultures with 50 μg/ml ascorbic acid for additional 10 days. Cocultures were then supplemented with 25 μM AT56 and DMSO as control for additional 7 days.

### RNA extraction and gene expression analyses

Cell cultures and sciatic nerves were homogenized in TriPure Reagent (Roche) using Precellys Lysing Kit CK14 and homogenizer (8000 rpm, 60 sec at 0°C, twice). RNA extraction was performed according to manufacturer’s instruction.

For RNAseq analyses, we prepared RNA samples from rat Schwann cell–mouse DRG neuronal co-cultures treated with 25 μM AT56 and DMSO as control (three different biological replicates/condition). Samples were processed at the Biotecnology Department at University of Verona as follows. Total RNA yield was determined using NanoDrop spectrophotometer (Thermo Fisher Scientific). Integrity of RNA samples was assessed using RNA 6000 Nano Kit (Agilent Technologies, Santa Clara, CA, USA) prior to library preparation. All samples showed an RNA integrity number (RIN) >8. RNAseq libraries were prepared from 2500 ng total RNA using the TruSeq RNA Library Preparation Kit v2 (Illumina, San Diego, CA, USA) after poly(A) capture, according to manufacturer’s instructions. Quality and size of RNAseq libraries were assessed by capillary electrophoretic analysis with the Agilent 4200 Tape station (Agilent). Libraries were quantified by real–time PCR against a standard curve with the KAPA Library Quantification Kit (KapaBiosystems, Wilmington, MA, USA). Libraries were then pooled at equimolar concentration and sequenced in 150PE mode on an Illumina NextSeq500 (Illumina). On average 42 million fragments were produced for each sample. Bioinformatic analysis was performed as follows. RNA sequences were aligned separately on mouse and rat deposited exome transcriptome sequences available at NCBI public database (mouse assembly GCF_000001635.24 and rat assembly GCF_000001895.5). We calculated relative gene expression in AT–56 treated cocultures versus DMSO treated by DESeq2 ^47^ and selected genes with a p–value < 0.001. We then classified as upregulated differentially expressed genes with fold change (FC) > 1.7 and as down regulated genes with FC < 0.5.

From mouse alignments, we identified 18 upregulated and 12 downregulated genes while from rat alignments we identified 24 upregulated and 2 downregulated genes. 300 ng of RNA were reversed transcribed using SSIV Superscipt (Invitrogen) following manufacturer’s instructions. To assess genes’ expression specifically in Schwann cells, we analyzed RNA total extract from rat Schwann cells–mouse DRG neuronal cocultures using primers pairs specific to rat mRNA sequences. Primers sequences were as follows: *Hmgcs2* forward 5’–CGC AGT CTA CCC AAG TGG TA–3’ and reverse 5’–GTC ATC GAG GGT GAA AGG CT–3’, *Acsl3* forward 5’–AGC AAA GGA GAC ACA TCC GTT–3’ and reverse 5’–CAG ATA TTC ATG AAT CGC TGT GTC–3’, *Pdk4* forward 5’–AAA CCG CCC TTT CCT GAC A–3’ and reverse 5’–GTC CCA TAG CCT GACATG GAA–3’, *Angptl4* forward 5’–ACC TTA AGA TAT GGC TGT TTT CTG CTG A–3’ and reverse 5’–CTG GGA ACC CTA TCT CCA GTC G–3’, *Gapdh* forward 5’–GGT TAC CAG GGC TGC CTT CTC TTG TGA–3’ and reverse 5’–CGG AAG GGG CGG AGA TGA TGA CCC T–3’, TBP forward 5’-CCT GTT CAG AAC ACC AAT AGT TTA–3’ and reverse 5’–GTG GAT ACA ATA TTT TGG AGC TGT–3’ Real Time–PCR (qRT–PCR) was performed using PowerUp SybrGreen master mix (Applied Biosystems), using 0.25 μM each primer and 5–15 ng cDNA for each reaction in triplicate. Cycling conditions were: 2 min at 50°c, 2 min at 95°C followed by 39 cycles of 30 sec 95°C, 30 sec annealing temperature, 1 min 30 sec at 72°C. Annealing temperatures were 60°C (*Acsl3*, *Hmgcs2* and *TBP*), 61°C (*Pdk4*), 62°C (*Angptl4*) and 64°C (*Gapdh*). Results were analyzed using the StepOne software (Applied Biosystems) according to manufacturer’s instructions.

Primers aligning on the mouse genome were used to detect the following transcripts: *Mbp*, *Mpz, l– pgds*, *gapdh, Hmgcs2*, *cpt1A*, *pdk4* and *36B4*. qRT–PCR analyses were performed using Evagreen master mix (Biorad), using 0.5 μM each primer and 15 ng cDNA for each reaction in triplicate. Cycling conditions were: 5 min at 95°C followed by 39 cycles of 30 sec 95°C, 30 sec at 60°C, 1 min at 72°C. Results were analyzed using the CFX Manager Software^TM^ from Biorad according to manufacturer’s instructions. Primers sequences were as follows: *Mbp* forward 5’–ACA CAC GAG AAC TAC CCA TTA TGG–3’and reverse 5’–GTT CGA GGT GTC ACA ATG TTC TTG–3’, *Mpz* forward 5’–CAC AAC CTA GAC TAC AGT GAC AAC G–3’ ‘and reverse 5’–TTC GAG GAG TCC TTC GAA GAT TTG–3’, *l–pgds* forward 5’–GGA GAA GAA AGC TGT ATT GTA TAT GTG C–3’ and reverse 5’– TAA AGG TGG TGA ATT TCT CCT TCA G–3’, *gapdh* forward 5’–TCA CCA GGG CTG CCA TTT GCA GTG G–3’ and reverse 5’–CGG AAG GGG CGG AGA TGA TGA CCC T–3’, *Hmgcs2* forward 5’-CAG AAT CAG TGG AAG CAA GCT G–3’ and reverse 5’–CAG AGT GGT GAG AGA GAA GTGAG–3’, *cpt1A* forward: 5’ AGG CCA CTG ATG ATG AAG GAG GG 3’ and reverse: 5’ GTT TGA GTT CCTCAC GGT CTA CC 3’, *pdk4* forward: 5’ GAA AAC CGT CCT TCC TTG ACC 3’ and reverse: 5’GTC TGT CCC ATA ACC TGA CAT AGA 3’, *36B4* forward 5’–AGA TTC GGG ATA TGC TGT TGG–3’and reverse 5’–AAA GCC TGG AAG AAG GAG GTC–3’.

### Immunofluorescence analyses

Schwann cell–DRG neuronal cocultures treated with Arachidonic acid or with 25 μM AT–56 (Cayman Chemicals) and 50 μM Etomoxir (Sigma Aldrich) were fixed in 4% PFA, permeabilized in 100% methanol at −20 °C for 15 min, blocked in 5% BSA (Sigma Aldrich), 1% donkey serum (Jackson Immuno Research), 0.2% Triton–X (Sigma) in PBS 1X (Gibco).

All antibodies were previously validated for the applications used. Primary antibodies used in immunofluorescence studies were: rat anti MBP hybridoma (diluted 1:2), chicken anti neurofilament M antibody (Covance PKC–593P, 1:1000), mouse anti Neurofilament SMI32 (BioLegend Cat# 801701, RRID:AB_2315331, 1:1000), rabbit anti turbo GFP (Thermo Fisher Scientific Cat# PA5–22688 Lot# UG2799941 RRID:AB_2540616, 1:500). Secondary antibodies were: rat anti Alexa Fluor 555 (Thermo Fisher Scientific Cat# A-21434, Lot#1987272 RRID:AB_2535855 1:1000), chicken anti Alexa Fluor 488 (Thermo Fisher Scientific Cat# A-11039, Lot#2304258, RRID:AB_2534096, 1:1000) and Rat Alexa fluor 647 (Thermo Fisher Scientific Cat# A21247, Lot#1921562, RRID AB_14778, 1:1000). Nuclei were stained by Hoechst. Tunel assay (Promega) was performed following manufacturer’s instructions before staining for MBP and NF. Cocultures treated with DNAse I were used as assay positive control. Slides were examined by epifluorescence by confocal microscopy on a Leica SP5.

### Preparation of detergent lysates and immunoblotting

All sciatic nerves from were lysed using the Precellys system (Bertin Instruments) in lysis buffer 2% (wt\vol) SDS, 25 mM Tris pH 7.4, 95 mM NaCl, 10 mM EDTA (all by Sigma), phosphatase inhibitor (PhoStop, Roche) and protease inhibitor (Complete Mini–EDTA free, Roche). Lysates from myelinated Schwann cell–neuronal cocultures were prepared and processed as described in ^16^. All lysates were boiled for 5 min @ 100°C and centrifuged for 10 min at 16,800 *g* at 16 °C. Supernatants were separated and proteins were quantified using Pierce BCA protein assay (ThermoFisher Scientific) according to manufacturer’s instructions.

Samples (8–25 μg protein extract) were run on 8–10% acrylamide gel at 100 Volt in Running Buffer (1X Tris–Glycine-SDS, Biorad) together with Precision Plus Protein Dual Color Standards (Biorad). Proteins were blotted on nitrocellulose membrane at 100 Volt at 4°C. Membranes were blocked in 5% milk in PBST (0.05% Tween in PBS 1X) and incubated for 16 hours at 4°C with the following primary antibodies: mouse anti–MBP (Covance Research Products Inc Cat# SMI–94R–100 Lot# RRID:AB_510039 and Cat# SMI–99P–100 Lot# RRID:AB_10120129, 1:4.000), mouse anti–NF (SMI 31 (BioLegend Cat# 801601, RRID:AB_10122491, 1:1000) chicken anti–MPZ (Millipore Cat# AB9352 RRID:AB_571090, 1:500), mouse anti–β–Tubulin (Sigma-Aldrich Cat# T4026, RRID:AB_477577), rabbit anti–Calnexin (Sigma Aldrich, Cat# C4731, RRID:AB_476845, 1:2000). Membranes were incubated with secondary antibodies all used 1:10000 and included anti–mouse IRdye 680 (LI–COR Biosciences Cat# 926–68070, RRID:AB_10956588) and anti–chicken IRdye 800 (LI–COR Biosciences Cat# 926–32218, RRID:AB_1850023). Quantitative Western Blotting analyses were performed using the Odyssey Infrared Imaging System (LI–COR Biosciences) according to the manufacturer’s instructions. We used the integrated intensity of the fluorescent signal to quantify protein expression of each sample using ImageJ software. Samples were normalized for the housekeeping protein tubulin or vinculin.

### Lipidomics and Metabolomics analyses

Lipidomics and metabolomics data were obtained by liquid chromatography coupled to tandem mass spectrometry. We used an API–4000 triple quadrupole mass spectrometer (AB Sciex, Farmington, MA, US) coupled to a HPLC system (Agilent) and CTC PAL HTS autosampler (PAL System) for lipidomic analysis. Metabolomics was performed on an API–3500 triple quadrupole mass spectrometer (AB Sciex, Farmington, MA, US) coupled with an ExionLC™ AC System (AB Sciex, Farmington, MA, US).

### Samples preparation

For lipidomics, sciatic nerves or Schwann cell–DRG neuronal cocultures were homogenised by tissue lyser for 2 min in 250 µl of ice–cold methanol/acetonitrile 50:50. Lysates were spun at 20,000g for 5 min at 4°C and supernatants passed through a 4 mm regenerated cellulose filter (200 nm Ø). Samples were finally dried under nitrogen flow at 40°C and resuspended in 100 μl in methanol/acetonitrile 50:50.

For metabolomics, sciatic nerves or cocultures homogenates were prepared by tissue lyser disruption for 2 min in 250 µl of ice-cold methanol/water/acetonitrile 55:25:20 containing [U–^13^C_6_]–glucose (Sigma Aldrich) 1 ng/µl and [U–^13^C_5_]–glutamine (Sigma Aldrich) 1 ng/µl as internal standards (IS). Lysates were spun at 15,000g for 5 min at 4°C and supernatants were passed through a 4 mm regenerated cellulose filter (4 mm Ø, Sartorius). Samples were then dried under nitrogen flow at 40°C and resuspended in 125 µl of methanol/water 70:30 for subsequent analyses.

Schwann cell–DRG neuronal cocultures were lysed in methanol/acetonitrile 50:50. LC–MS/MS runs were performed on an API–4000 triple quadrupole mass spectrometer (AB Sciex, Farmington, MA, US) coupled to a HPLC system (Agilent) and CTC PAL HTS autosampler (PAL System). Results were obtained after correction for natural abundance of ^13^C and expressed as Mass Isotopomer Distribution (MID). MultiQuant™ software (version 3.0.3, AB Sciex, Farmington, MA, US) was used for data analysis and peak review of chromatograms.

All results were normalized over the total protein content as determined by BCA assay on protein fractions after metabolites/lipids extraction. *In vivo* analyses were performed on sciatic nerves from at least 4 *l–pgds^-/-^* and wild type littermate control mice, at 4, 6 and 8 months and on sciatic nerves from at least 3 *P0–Cre//l–pgds^flx/flx^* and littermates’ control (*l–pgds^flx/flx^*) mice at 8 and 14 months. *In vitro* analyses were performed on at least 3 different biological samples of Schwann cell–DRG neuronal cocultures.

### Phospholipids

Phospholipids were identified and evaluated by LC–MS/MS on an API–4000 triple quadrupole mass spectrometer (AB Sciex) coupled to a HPLC system (Agilent) and CTC PAL HTS autosampler (PAL System). Methanolic extracts were analyzed by using a XTerra Reverse Phase C18 column (3.5 μm 4.6X100mm, Waters) and Methanol (MetOH) with 0.1% formic acid as isocratic mobile phase for positive ion mode in 5 min total run for each sample. Negative ion mode analysis was conducted with a cyano–phase LUNA column (50 mm X 4.6 mm, 5 μm; Phenomenex) and 5 mM ammonium acetate pH 7 in MetOH as isocratic mobile phase in 5 min total run for each sample. The identity of the different phospholipid families was confirmed using pure standards, namely one for each family and lipidomics was quantified by external standard method. Collectively, we analysed more than 200 phospholipids belonging to following classes: phosphatidic acids (PA), lysophosphatidic acid (LPA) phosphatidylcholines (PC), lysophosphatidylcholines (LPC), phosphatidylethanolamines (PE), lysophosphatidylethanolamines (LPE), phosphatidylserines (PS), phosphatidylinositols (PI), lysophosphatidylinositols (LPI) phosphatidylglicerols (PG), sulfatides (Sul), ceramides (Cer), Galactosyl ceramide (Gal Cer) and sphingomyelins (SM). Free and total cholesterol levels were determined in sciatic nerves methanolic extracts by cholesterol quantitation kit (MAK043–1KT–Sigma Aldrich) following manufacturer’s instructions.

### Metabolites

Quantification of energy metabolites (glycolysis, pentose phosphate pathway and Krebs cycle intermediates) was performed using a cyano–phase LUNA column (50 mm x 4.6mm, 5 µm; Phenomenex) by a 5 min run in negative ion mode. We used water as stationary phase (A) and 2 mM ammonium acetate in MetOH as mobile phase (B). 10% A and 90% B gradient and flow rate of 500 µl/min were applied for all analyses. Carnitine quantification was performed on acetonitrile/methanol extracts by using a Varian Pursuit XRs Ultra 2.8 Diphenyl column. Samples were analysed by a 3 min run in positive ion mode using 0.1% formic acid in MetOH as mobile phase. All metabolites analysed in the described protocols were previously validated by pure standards and quantified by standard curves and different internal standards ^48, 49^.

### Amino Acids

Amino acids, their derivatives and biogenic amine quantification was performed through previous derivatization. Briefly, 25µl out of 125µl of samples were collected and dried separately under N2 flow at 40°C. Dried samples were resuspended in 50µl of phenyl-isothiocyanate (PITC), EtOH, pyridine and water 5%:31.5%:31.5%:31.5% and then incubated for 20 min at RT, dried under N2 flow at 40°C for 90 min and finally resuspended in 100µl of 5mM ammonium acetate in MeOH/H2O 50:50. Quantification of different amino acids was performed by using a C18 column (Biocrates, Innsbruck, Austria) maintained at 50°C. The mobile phases for positive ion mode analysis were phase A: 0.2% formic acid in water and phase B: 0.2% formic acid in acetonitrile. The gradient was T0: 100%A, T5.5: 5%A, T7: 100%A with a flow rate of 500µl/min. All metabolites analyzed in the described protocols were previously validated by pure standards and internal standards were used to check instrument sensitivity.

### Metabolites flux analyses

MultiQuant™ software (version 3.0.3, AB Sciex, Farmington, MA, US) was used for data analysis and peak review of chromatograms. Raw areas were normalized by the median of all metabolite areas in the same sample. Specifically, we defined the relative metabolite abundance (m_a_^N^) as:

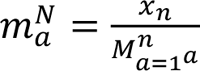

where *x_n_* represents the peak area of metabolite *n* for samples *a, b, …, z*, and *M^n^_a=1^a^_* represents the median of peak areas of metabolite *n* for samples *a, b, …, z*. Obtained data were then transformed by generalized log-transformation and Pareto scaled to correct for heteroscedasticity, reduce the skewness of the data, and reduce mask effects ^50^. In detail, obtained values were transformed by log10 and obtained values underwent Pareto scaling as follows:

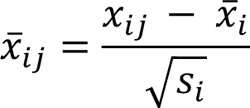

where *x_ij_* is the transformed value in the data matrix (*i* (metabolites), *j* (samples)) and *s_i_* is the standard deviation of transformed metabolite values ^51^. Obtained values were considered as relative metabolite levels. Data processing and analysis were performed by MetaboAnalyst 5.0 web tool ^52^.

### Acetate and Ketone bodies quantitation

Acetate’s content was determined by Ketone Body Assay Kit (MAK134 Sigma Aldrich). Schwann cell–DRG neuronal cocultures and nerves were homogenized using the Precellys System and tested for Acetoacetate content following manufacturer’s instructions. β–Hydroxybutyrate (BOH) content was assessed in coculture media supernatant and in sciatic nerves, following manufacturer’s instructions. Results were normalized over the total protein content as determined by Pierce BCA assay.

### Statistical analyses

Data were collected randomly and assessed blindly on samples of comparable size, especially for all *in vitro* analyses. The data distribution was assumed to be normal, although we did not formally test it. All statistical analyses were performed on at least three different experiments. Statistical detailed analyses are reported in each figure legends and all assays were performed using the Prism 9 Software package (GraphPad). A method checklist is available with the supplementary materials

### Data availability

The datasets generated during and/or analyzed during the current study are available from the corresponding author on reasonable request.

## ACKNOWLEDGEMENTS

We are in debt to M. Delledonne and M. Rossato (University of Verona and Personal Genomics) for performing the transcriptomic analyses and P. Canevazzi (IRCCS, San Raffaele Scientific Institute) for morphological analyses. We are also grateful to Yoshihiro Urade (Tsukuba University, Japan) for kindly providing *l–pgds* and *l-pgds^flx/flx^* mutant mice and V. Lee (Perelman School of Medicine at the University of Pennsylvania) for the MBP antibody. The model shown in the manuscript was created with BioRender.com. This study was supported by National Institute of Health/National Institute of Neurological Disorders and Stroke (R01NS099102) (C.T.) and partially by the Ministry of University and Research (MUR) Progetto Eccellenza (2018–2022) to the Department of Pharmacological and Biomolecular Sciences, DiSFeB, Università degli Studi di Milano (M.A., D.C. and N.M.).

## AUTHOR CONTRIBUTION

A.T. designed the experimental plan, conducted the majority of the experiments and wrote the manuscript. R.L.M, M.C. and M.F. contributed to *in vitro* and *in vivo* studies. M.A, S.P., G.I., D.C. and N.M. performed metabolomics and lipidomic studies, contributed to the experiment design and helped in writing the manuscript. A.C. and L.M. performed bioinformatic studies. P.P., G.D. and A.Q. performed morphological analyses. C.T. designed the experimental plan, supervised the project and wrote the manuscript. All authors commented on the manuscript.

## SUPPLEMENTARY FIGURES

**Supplementary Figure 1.** In vivo lipidomic and metabolomics profiles. Heat–map diagrams showing the top 50 dysregulated metabolites detected and quantified in wild type littermate controls (WT) and *l–pgds^-/-^* (KO) sciatic nerves at 4 months (**a**), 6 months (**b**) and 8 months (**c**). Heat–map diagram showing the top 50 dysregulated metabolites detected and quantified in wild type littermate controls (WT) and in *P0–Cre//l–pgds^flx/flx^*sciatic nerves at 8 (**d**) and 14 (**e**) months. Acronyms’ legend: (PA), lysophosphatidic acid (LPA) phosphatidylcholines (PC), lysophosphatidylcholines (lysoPC), phosphatidylethanolamines (PE), lysophosphatidylethanolamines (lysoPE), phosphatidylserines (PS), phosphatidylinositols (PI), lysophosphatidylinositols (LPI) phosphatidylglicerols (PG), sulfatides (Sul), ceramides (Cer), Galactosyl ceramide (Gal Cer) and sphingomyelins (SM), PEP: phosphoenolpyruvate, αKG: alpha–ketoglutarate, OAA: oxalacetate.

**Supplementary Figure 2.** Morphological analyses of *Chat–Cre//l–pgds^flx/flx^* nerves. **a)** Genotyping PCR for *l–pgds* and for *Chat–Cre* alleles on genomic DNA prepared from P7 ventral (V) and dorsal (D) spinal cord regions of *Chat–CRE//l–pgds^flx/flx^* mutants and littermate controls. The 444 bp *l-pgds* null allele (*) is present only in the ventral spinal cord region of *l–pgds^flx/flx^* mice expressing the Cre recombinase. The *l–pgds^flx/flx^* allele (396 bp, **) is present in all spinal cord samples. To validate the results we included control samples of genomic DNA prepared from tails of *l–pgds^flx/flx^* (f/f), wild type (+/+) and heterozygous *l–pgds* (+/–) mice. The wild type allele (290 bp, ***) is present only in wild type (+/+) and heterozygous *l–pgds* (+/–) genomic DNA. **b)** Electron microscopy analyses of *Chat–CRE//l–pgds^flx/flx^* and littermate controls injured sciatic nerves at P7. Scale bar: 2 µm. **c)** Graph showing *g* ratios as a function of axon diameter in P7 *Chat–CRE// l–pgds^flx/flx^* nerves (red line) as compared to littermate controls (black line). (g ratio WT: 0.702 ± 0.0036; *Chat–CRE// l–pgds^flx/flx^*: 0.723 ± 0.0036. Unpaired t test analyses; *P = 0.0107, t = 3.968, df = 5). N = 3 control mice; N = 4 *Chat– CRE// l–pgds^flx/flx^* mice. **d)** Electron microscopy analyses of *Chat–CRE//l–pgds^flx/flx^* and littermate controls injured sciatic nerves at P30. Scale bar: 2 µm. **e)** Graph showing *g* ratios as a function of axon diameter in P30 *Chat–CRE// l–pgds^flx/flx^* nerves (red line) as compared to littermate controls (black line). (g ratio WT: 0.671 ± 0.0068; *Chat–CRE// l–pgds^flx/flx^*: 0.682 ± 0.0074. Unpaired t test analyses; P = 0.324 n.s., t = 1.124, df = 4). N = 3 mice/genotype. **f)** Semi–thin sections of 8 months wild–type littermate controls (WT) and *Chat–CRE// l–pgds^flx/flx^* sciatic nerves (KO). Bar: 50 μm. **b**) Graph, average of three different experiments, showing the percentage of myelin morphological aberrations in 8 months old wild type littermate controls (white bar) and *Chat–CRE// l–pgds^flx/flx^*sciatic nerves (red bar). Alterations were determined as the number of fibers presenting myelin structural alterations (pink arrows) over the total number of fibers in the entire reconstructed nerve cross section. Error bars represent mean ± s.e.m. N = 3 different mice/genotype. (Unpaired t test analysis, P = 0.8806 n.s.; t = 0.160 df = 4).

**Supplementary Figure 3.** Developmental myelination is normal in *P0–Cre//l–pgds^flx/flx^* nerves. **a)** Electron microscopy analyses of *P0–CRE//l–pgds^flx/flx^* and littermate controls injured sciatic nerves at P7. Scale bar: 5 µm. **b)** Graph showing *g* ratios as a function of axon diameter in P7 *P0–CRE//l–pgds^flx/flx^* nerves (red line) as compared to littermate controls (black line). (g ratio WT: 0.6986 ± 0.0139; *P0–CRE//l–pgds^flx/flx^*: 0.7054 ± 0.0123. Unpaired t test analyses; P = 0.7343 n.s., t = 0.364, df = 4). N = 3 mice/genotype. **c)** Distribution of myelinated fibers is similar in 7 days *P0–CRE//l–pgds^flx/flx^*nerves and littermate controls. (Fisher’s exact test; P = 0.7379 (total versus 1–1.5 μm), P = 0.4094 (total versus 1.5–2 μm), P = 0.164 (total versus 2–2.5 μm), P = 0.1122 (total versus 2.5–3 μm), P = 0.4471 (total versus 3–3.5 μm)). Over 40 fibers for each genotype were counted. N = 3 mice/genotype. **d)** Electron microscopy analyses of *P0–CRE//l–pgds^flx/flx^* and littermate controls injured sciatic nerves at P30. Scale bar: 5 µm. **e)** Graph showing *g* ratios as a function of axon diameter in P30 *P0–CRE//l–pgds^flx/flx^*nerves (red line) as compared to littermate controls (black line). (g ratio WT: 0.6524 ± 0.0093; *P0–CRE// l–pgds^flx/flx^*: 0.659 ± 0.0024. Unpaired t test analyses; P = 0.5306 n.s., t = 0.6857, df = 4). N = 3 mice/genotype. **f)** Distribution of myelinated fibers is similar in 30 days *P0–CRE//l–pgds^flx/flx^* nerves and littermate controls. (Fisher’s exact test; P = 0.3402 (total versus 1–1.5 μm), P = 0.8216 (total versus 1.5–2 μm), P = 0.4637 (total versus 2–2.5 μm), P = 0.8875 (total versus 2.5–3 μm), P = 0.5962 (total versus 3–3.5 μm), P = 0.2178 (total versus >3.5 μm)). Over 50 fibers for each genotype were counted. N = 3 mice/genotype. **g)** Semi–thin sections of 10 months wild–type littermate controls (WT) and *P0–CRE// l–pgds^flx/flx^* sciatic nerves (KO). Bar: 50 μm. **h)** Graph showing *g* ratios as a function of axon diameter in 10 months *P0–CRE//l–pgds^flx/flx^* nerves (red line) as compared to littermate controls (black line). (g ratio WT: 0.6631 ± 0.0058; *P0–CRE//l–pgds^flx/flx^*: 0.6685 ± 0.009. Unpaired t test analyses; P = 0.6499 n.s., t = 0.4898, df = 4). N = 3 mice/genotype. **i)** Distribution of myelinated fibers is similar in 10 months *P0–CRE//l–pgds^flx/flx^* nerves and littermate controls. (Fisher’s exact test; P = 0.9267 (total versus 1–2 μm), P = 0.7465 (total versus 2–3 μm), P = 0.1166 (total versus 3–4 μm), P = 0.246 (total versus 4–6 μm)). Over 250 fibers for each genotype were counted. N = 3 mice/genotype.

**Supplementary Figure 4.** Main dysregulated lipids and metabolites identified in *l–pgds^-/-^* mutants. **a)** Scheme showing the omega-6 fatty acid inter–conversions catalyzed by Δ6-desaturase and elongase enzymes. **b–d**) Graph showing the differences in lysophosphatydilcholines contents in wild type littermate controls (WT) and *l–pgds^-/-^* (KO) sciatic nerves at 4, 6 and 8 months. Values are relative to the analyses performed in Fig. 3. Variation in lipids’ amount is expressed as fold change to WT arbitrarily set as 1.0. Error bars represent mean ± s.e.m. (Unpaired t-test. LPC 18:2 6 months ***P = 0.0007, t=4.867, df =10; LPC 18:2 8 months **P = 0.0098, t=3.178, df =10; LPC 20:3 6 months **** P <0.0001, t=8.214, df =10; LPC 20:3 8 months *P = 0.0197, t=3.38, df =5; LPC 20:4 6 months **** P <0.0001, t=8.631, df =10; LPC 20:3 8 months *P = 0.0314, t=2.962, df =5). **e)** Graph showing the differences in acetyl–CoA contents in wild type littermate controls (WT) and *l– pgds^-/-^* (KO) sciatic nerves at 4, 6 and 8 months. Values are relative to the analyses performed in Fig. 4. Error bars represent mean ± s.e.m. (Unpaired t-test. 8 months *P = 0.0367, t=2.828, df =5). **f)** Graph showing the differences in citrate contents in wild type littermate controls (WT) and *l–pgds^-/-^*(KO) sciatic nerves at 4, 6 and 8 months. Values are relative to the analyses performed in Fig. 4. Error bars represent mean ± s.e.m. (Unpaired t-test. 6 months **P = 0.0047, t=3.614, df =10; 8 months *P = 0.038, t=2.799, df =5). **g)** Graph showing the differences in α–ketoglutarate contents in wild type littermate controls (WT) and *l–pgds^-/-^* (KO) sciatic nerves at 4, 6 and 8 months. Values are relative to the analyses performed in Fig. 4. Error bars represent mean ± s.e.m. (Unpaired t-test. 8 months *P = 0.0473, t=2.617, df =5). **h)** Graph showing the differences in lactate contents in wild type littermate controls (WT) and *l–pgds^-/-^* (KO) sciatic nerves at 4, 6 and 8 months. Values are relative to the analyses performed in Fig. 4. Error bars represent mean ± s.e.m. (Unpaired t-test. 6 months **P = 0.0066, t=3.418, df =10).

**Supplementary Figure 5.** L–PGDS is not implicated in *in vitro* neuronal and Schwann cell survival. **a)** Representative immunofluorescence coupled to Tunel assay of mouse DRG – rat Schwann cells myelinated cocultures. After 14 days in myelinating conditions, cultures were treated for additional 7 days with 25 μM AT–56 or with DMSO as control vehicle. At the end of the treatment cultures were fixed and stained for MBP (white) and processed for TUNEL assay. Control cocultures were treated with 10 u/ml DNase1 as positive control for Tunel assay. AT–56 treatments in already myelinated cultures induces myelin degeneration, but not cell death. N = 3 different independent coculture experiments. Bar: 100 μm. **b)** Representative immunofluorescence coupled to Tunel assay of mouse DRG – rat Schwann cells myelinated cocultures treated with 300 μM Arachidonic acid (complexed to BSA) for additional 7 days after 14 days in myelinating conditions. At the end of the treatment cultures were fixed and stained for MBP (white) and processed for TUNEL assay. Control cocultures were treated with 10 u/ml DNase1 as positive control for Tunel assay. Myelin degeneration was observed upon 300 μM Arachidonic acid treatment with no signs of Schwann cell death. The same amount of BSA was added to control cocultures. N = 3 different independent coculture experiments. Bar: 100 μm.

**Supplementary Figure 6.** Ketone bodies synthesis is critical to maintain the integrity of the axo-glial unit. **a)** qRT–PCR analyses on mRNA prepared from myelinated wild type Schwann cell – neuronal cocultures infected with shRNA lentiviruses against *Hmgcs2* (*sh062*, *sh362*, *sh372*) or scramble sh RNA (*shscr*). Three days post infection, cocultures were supplemented with ascorbic acid to allow myelination for 10 days, then treated with 25 uM AT–56 or vehicle only, as control, for additional 7 days. *Hmgcs2* expression is significantly reduced specifically in AT–56 treated cocultures infected with *Hmgcs2* targeting lentiviruses. Data have been normalized to *36B4* expression level and analyzed with the StepOne Software v2.3 (Applied Biosystems). N = 3 different mRNA preparations and analyses. Error bars represent mean ± s.e.m. (Unpaired t test; *shscr* AT56–*sh062AT–56* ***P = 0.0001 (t=14.33; df=4); *shscr* AT56–*sh362AT–56* ***P = 0.0002 (t=13.81; df=4); *shscr* AT56–*sh372AT–56* ****P < 0.0001 F (t=18.56; df=4). **b)** Representative immunofluorescence of wild–type myelinated cocultures Schwann cells neuronal cocultures infected with *Hmgcs2 sh362* or *Hmgcs2 sh062*. Three days post infection, cocultures were supplemented with ascorbic acid to allow myelination for 10 days, then treated with 25 uM AT–56 or vehicle only, as control, for additional 7 days. At the end of the experiments cultures were fixed and stained for GFP to visualize infected cells (fluorescein), non–phosphorylated neurofilament SMI–32 (rhodamine) and MBP (white). Ablation of *Hmgcs2* coupled to *L-pgds* block, causes axonal instability. *N* = 3 independent biologic replicates. Bar: 100 μm.

**Supplementary Figure 7.** Schematic representation of Schwann cell metabolic rewiring occurring during myelin degeneration. Loss of L–PGDS enzymatic activity determines the release of Arachidonic Acid (AA) and fatty acids from myelin sheath lysophosphatidylcholines (LPC). Free fatty acids enter the mitochondria via *Acsl3* and *Cpt1a*, where β–oxidation convert them into Acetyl–CoA (Ac–CoA). Next, *Hmgcs2* drives synthesis of ketone bodies, acetoacetate (AcAc) and β–hydroxybutyrate (BOH), from accumulating Ac–CoA. While BOH exits Schwann cells and it is metabolized in the axon, acetoacetate sustains the TCA cycle in Schwann cells. In parallel, Ac–CoA production deriving from glucose is hampered by upregulated expression of *Pdk4*. Created with BioRender.com.

